# Co-optation of Transcription Factors Drives Evolution of Quantitative Disease Resistance Against a Necrotrophic Pathogen

**DOI:** 10.1101/2025.02.26.640262

**Authors:** S. Einspanier, C. Tominello-Ramirez, F. Delplace, R. Stam

## Abstract

Wild relatives of crop species possess diverse levels of quantitative disease resistance (QDR) to biotic stresses, yet the genomic and regulatory mechanisms underlying these differences are poorly understood. In particular, how QDR against a generalist necrotrophic pathogen evolved and whether it is driven by conserved or species-specific regulatory networks remains unclear. Here, we examined the transcriptomic responses of five diverse wild tomato species that span a gradient of QDR. We initially hypothesised that conserved regulatory modules might control QDR. Instead, we use differential gene expression analysis and weighted gene co-expression network analysis (WGCNA) to find that species-specific regulatory features, encompassing both infection-induced and constitutively expressed genes, predominantly shape QDR levels. Although we identified an ethylene response factor among candidate genes for QDR-regulation, it did not fully account for the phenotypic variation. To further dissect the evolutionary basis of these regulatory patterns, we performed phylotranscriptomic analyses on gene regulatory networks. Notably, our findings reveal that the conserved NAC transcription factor 29 is pivotal in developing disease resistance only in *S. pennellii*. The differential regulation and altered downstream signalling pathways of NAC29 provide evidence for its co-option in the resistance mechanisms of *S. pennellii*. This finding highlights the species-specific rewiring of gene regulatory networks by repurposing a conserved regulatory element to enhance resistance against pathogens effectively. These results offer new insights into the evolutionary and regulatory complexity underlying QDR and emphasise the significance of species-specific gene regulation in shaping resistance against a cosmopolitan necrotrophic pathogen.

## Introduction

Developing or engineering durable resistance against necrotrophic pathogens in crop plants displays a major bottleneck in modern plant breeding. Although the scientific community gained a comprehensive understanding of dominant R-gene-mediated resistance, especially against biotrophic pathogens, an R-gene-mediated resistance against necrotrophic generalist pathogens like *Sclerotinia sclerotiorum* or *Botrytis cinerea* has not yet been described (Mbengue et al. 2016; Wang et al. 2019). Accordingly, the primary goal of resistance breeding against necrotrophic pathogens is to enhance quantitative disease resistance (QDR), where many independent loci contribute marginally to a continuous pattern of resistance (Caseys et al. 2021). The current state of knowledge about mechanisms involved in QDR has been reviewed by (Corwin and Kliebenstein 2017; Poland et al. 2009; Roux et al. 2014; Gou et al. 2023).

Natural or wild plant pathosystems are a valuable tool for unravelling the complexity of the QDR mechanisms and their evolution, especially when looking into crop wild relatives (Kahlon and Stam 2021). Crop wild relatives harbour a wide diversity of morphological properties and represent a common source of novel traits in modern crops (Szymański et al. 2020; Albaladejo et al. 2017; Nosenko et al. 2016). Adapting to diverse habitats is accompanied by the evolution of resistance or tolerance against numerous stresses, including drought, temperature, and disease (Wei et al. 2024; Bolger et al. 2014). For instance, previous studies have demonstrated that accessions of the crop wild relative *Solanum chilense* exhibit a wide inter- and intrapopulation diversity in resistance response against pathogens like *Phytophthora infestans, Fusarium oxysporum* and *Alternaria* spp. (Kahlon et al. 2021; Stam et al. 2017; Schmey et al. 2023). Several wild tomato species also show distinct patterns of presence-absence variation, copy number variation, and specific patterns of selection on pathogen resistance genes within and between populations or species (Stam et al. 2019; Seong et al. 2020; Silva-Arias et al. 2025). We have previously shown that wild tomato species employ distinct mechanisms to achieve QDR against *S. sclerotiorum*, particularly by modulating the duration of the asymptomatic phase (lag phase) and the speed of lesion growth (lesion doubling time, LDT). These measures appear not correlated and exhibit significant host and population specificity. Thus, species or populations might have adapted specific strategies to regulate QDR. I.e. prolonging the lag phase in one case or increasing the pathogen’s LDT in another (Einspanier et al. 2024). The possibility of such differentiation is supported by population genomic analyses, showing distinct genomic separation and differentiation among *S. chilense* populations, which also exhibit diversifying elicitor responses and phytohormonal regulation (Kahlon et al. 2023; Stam et al. 2017; Böndel et al. 2015). Interestingly, a previous study investigating *S. pennellii* introgression lines described that genomic variation did not explain the phenotypic variability and ranked regulatory variability as the main determinant of *B. cinerea* resistance on fruits, highlighting how regulatory plasticity can enable new resistance traits (Szymański et al. 2020).

The underlying regulatory interplay governing QDR remains elusive. Although recent studies identified QDR-associated quantitative trait loci, the explained phenotypic variation remains relatively low (Thomas et al. 2024; Pink et al. 2022). While the genomic characterisation of QDR requires many resources (such as mapping populations, multiparent advanced generation intercross populations or introgression lines of sufficient size, (Corwin and Kliebenstein 2017; Thomas et al. 2024), RNA sequencing is suitable for determining QDR regulation, leveraging high throughput and decreasing costs to support large-scale sampling. This enables researchers to investigate complex regulatory networks, thereby identifying and characterising nuanced shifts in gene expression and their evolution (Gómez-Picos and Eames 2015; Delplace et al. 2020).

Gene networks are a powerful tool for characterising relationships or interactions among genes and help understand the underlying molecular mechanisms of various phenotypes (Delplace et al. 2022; Ovens et al. 2021). Typically, gene network analyses are separated into undirected coexpression networks (like weighted gene correlation network analysis, WGCNA) and directed gene regulatory networks (GRNs, (Li et al. 2015; Langfelder and Horvath 2008). Albeit fundamentally different in the underlying statistical concepts, both GRNs and WGCNAs can be used to characterise complex regulatory networks, as previously shown in cultivated tomatoes. Tominello-Ramirez et al. (2024) integrated WGCNA and GRN to characterise a specific class of ethylene response factors involved in defence against necrotrophic fungi of the genus *Alternaria*, illustrating how a system approach can be facilitated to describe QDR networks and their moderators. Although certain gene networks appear to be highly conserved (with some predating the existence of land plants), there is increasing evidence that gene regulation network evolution can be a relatively quick response to different stresses (Curci et al. 2022; Wu et al. 2021; Obertello et al. 2015; Crow et al. 2022; Wei et al. 2024). Accordingly, evolutionary flexibility was hypothesised to enhance QDR robustness (Derbyshire and Raffaele 2023). However, it remains unclear how network evolution drives QDR and how phylogenic relationships determine the degree of QDR. This is mostly due to the high analytical complexity, challenging construction of cross-species networks and multidimensional comparative analysis on non-model organisms exhibiting a wide range of complex phenotypes (Schoenrock et al. 2017).

This also involves the development of QDR against the generalist pathogen *S. sclerotiorum*, which infects a broad range of hosts, including agronomically relevant crops such as oilseed rape, sunflower, and tomatoes (Derbyshire et al. 2022; Derbyshire and Raffaele 2023). Although Sucher et al. (2020) showed signs of recent gene expression acquisition and exaptation in a group of ATP Binding Cassette type G transporters responding to *S. sclerotiorum* infection; the regulatory cues driving enhanced resistance through network reconfiguration against this pathogen remain unknown.

Here, we examine transcriptome dynamics and plasticity to identify both conserved and newly recruited regulators of QDR-level. We employ a novel approach integrating phylotranscriptomic analysis and network inference to elucidate gene network evolution and unravel the regulatory cues driving QDR. Specifically, we investigate the regulatory architecture underlying QDR against a generalist necrotrophic pathogen, comparing the rewiring of gene regulatory networks in resistant versus susceptible genotypes across five tomato species (*S. chilense*, *S. habrochaites*, *S. lycopersicoides*, *S. pennellii*, and *S. pimpinellifolium*). We test the hypothesis that shared regulatory networks drive QDR while downstream mechanisms fine-tune species-specific regulatory responses. By integrating phylogenetic insights with transcriptomic data, we uncover patterns of gene family expansion, functional divergence, and regulatory remodelling that collectively enhance QDR. This evolutionary approach helps pinpoint the “ancient core” of plant immune systems shared across species and novel, rapidly evolving genes that can contribute to the partial, durable form of resistance in wild tomato relatives. Our findings offer a valuable resource that contrasts regulatory responses among genotypes with varying QDR levels, laying the groundwork for further exploration of the functional mechanisms shaping QDR.

## Materials and Methods

### Plant growth conditions

We obtained germplasm from the C. M. Rick Tomato Genetics Resource Center at UC Davis (see suppl. tab. 1, TGRC UC-Davis, https://tgrc.ucdavis.edu/) and cultivated the plants in the greenhouse of the Phytopathology Department at Christian Albrechts University Kiel, Germany. We surface-sterilized the seeds with a 2.75% hypochlorite solution. We propagated mature plants through cuttings using Chryzotop Grün 0.25% in Stender C700 substrate. We maintained the growing environment at approximately 21 °C (±10 °C), 65% relative humidity, and a 16-hour photoperiod. We fertilised the plants monthly using a drip irrigation system with a 1% Sagaphos Blue solution. For further methodological details, please refer to (Einspanier et al. 2024; Tominello-Ramirez et al. 2024).

### Fungal growth conditions

We freshly grew the fungus *Sclerotinia sclerotiorum* (1980) on potato dextrose agar (Sigma-Aldrich) at 25 °C in the dark. We performed the inoculation using a liquid mycelium macerate, following the method described in Einspanier et al. (2024). Briefly, we incubated 100 mL of potato dextrose broth with four 1 cm pieces of fully overgrown PDA on a rotary shaker (120 rpm, 24 °C) for four days. After incubation, we mixed the culture using a dispenser and vacuum-filtered it through cheesecloth. We then concentrated the filtrate to an optical density at 600nm (OD_600_) of 1 using the clear supernatant as dilution. We used fresh potato dextrose broth as a negative control and added Tween80 as a surfactant.

### Experimental conditions

The experimental conditions of the sequencing experiment were identical to previous experiments conducted to measure LDT (Einspanier et al. 2024). In short, we placed detached leaflets on wet tissue papers in a tray with the adaxial side up. Then, the leaves were inoculated with 10 µL of fungal mycelial mixture (OD_600_ =1) or empty PDB. The hood was covered and incubated at 23 °C. LED lights were used to continue the 16-h photoperiod. We sampled relative to the lesion development in the middle of lesion spread, but always at the same daytime, minimising the influence of the circadian rhythm on gene expression. A sterile scalpel was used to sample a 2cm x 2cm big piece covering the lesion and the surrounding tissue. Two leaf segments were pooled into one sample. We sampled four biological replicates per condition and genotype. The leaf segments were submerged in 750 µL DNA/RNA shield (Zymo Research) in ZR BashingBead Lysis Tubes (2 mm). We used the Zymo Research Quick-RNA Plant Kit for total RNA isolation following the manufacturer’s protocol. We used a NanoDrop One (ThermoFisher) and agarose gels to determine RNA quality and integrity. For RNA yield quantification, we used a Fluorometer (Promega Quantus).

### Sequencing and library preparation

We in-house prepped a Lexogen QuantSeq 3’ mRNA-Seq V2 (Lexogen, Vienna, Austria) library, following the manufacturer’s guidelines. Spike-RNA was added for quality assessment, as was the PCR Add-on Kit, to determine the correct number of PCR cycles required to amplify mRNA sequences. The library was sequenced at the Competence Centre for Genomic Analysis Kiel (CCGA) on an Illumina NovaSeq 600 (Illumina, San Diego, CA, USA) with 2 x 100 bp aiming for five mio. reads per sample.

### Bioinformatic preprocessing

We performed a quality assessment on the raw reads using FastQC/MultiQC (Simon Andrews 2010; Ewels et al. 2016). Then, we used Cutadapt v4.8 for initial filtering and adapter trimming following recommendations from the library manufacturer (Martin 2011). This included the settings `-a “polyA=A{20}” -a “QUALITY=G{20}”` and custom options for adapter removal (see online resources). Subsequently, we depleted ribosomal RNA bioinformatically using a custom pipeline (see online resources). For this, we generated reference-rRNA sequences for all five species based on 45S rDNA-sequences of the 5.8S, 18S and 25S subunits from the *A. thaliana* quality reference genome (GeneBankIDs: 5.8S: AB373816.1, 25S: OK073662.1 18S: OK073663.1, (Rabanal et al. 2017) and the chloroplast/mitochondrial rRNA of *S. lycopersicum* (GeneIDs: 34678306, 3950431, 3950467, 3950435, 3950433, 34678288, 34678318, 34678306). We then extracted species-specific rRNA sequences using Blast v2.13 (Camacho et al. 2009). We inflated the flanking sequences by 100 bp and extracted the final sequences using bedtools v2.31.1 ‘getfasta’ (Quinlan and Hall 2010). We used the following reference genomes: *S. lycopersicoides* (BioProject PRJNA727176, (Powell et al. 2022), *S. chilense* (BioProject: PRJNA1210999), *S. pennellii* (BioProject: PRJEB5809, (Bolger et al. 2014), *S. habrochaites* (BioProject: PRJCA008297,(Yu et al. 2022), and *S. pimpinellifolium* (BioProject: PRJNA607731, (Wang et al. 2020b). Trimmed reads of each species were mapped on the respective rRNA reference using the STAR-aligner v2.7.9 with custom settings retaining only non-mapping reads (see online resources, (Dobin et al. 2013). Then, we mapped the rRNA-depleted reads against the respective reference genome using the STAR aligner with settings, following the library manufacturer’s recommendations (see mapping statistics in suppl. tab. 2).

We used the RNAseq read-assisted tool GeneExt (https://github.com/sebepedroslab/GeneExt, last accessed Oct. 2024) to extend or predict missing UTR regions in our genome annotations. By using the options `--peak_perc 10 --orphan -j 16 -v 1 -m 5000`, we enhanced the annotation completeness significantly. (Zolotarov et al. 2023). Next, we quantified aligned sequencing reads with featureCounts v2.0.6 using custom options `-s 1 -T 16 -M -t exon -g gene_id` as this sequencing library allows only the quantification of expression on gene-level (Liao et al. 2014).

### Annotation and Proteome

We developed a custom pipeline to improve the proteome of the five host plant species. First, we utilised per-species GeneExt-curated genome annotations to extract transcripts using gffread v0.12.7 and extracted protein sequences based on the longest open reading frames (ORF) with TransDecoder v5.7.1 (Pertea and Pertea 2020; Haas 2024a, 2024b). To retain proteins with functional significance, we employed BLASTp to identify homologous proteins in the UniProt database using the parameters `-max_target_seqs 1 -evalue 1e-5.` (Camacho et al. 2009; UniProt Consortium 2017). We kept only those proteins with significant UniProt matches for downstream analysis. For protein sequences that did not achieve a sufficient match in UniProt, we further assessed their homology to the ITAG4 proteome of *S. lycopersicum* (https://solgenomics.net/organism/Solanum_lycopersicum/genome/) using Orthofinder to retain tomato-specific proteins. Next, we evaluated sequences with low identity to the tomato proteome by assigning functional annotations using PANNZER2 (Törönen and Holm 2022). We retained all sequences with a PANNZER2 positive predictive value (PPV) score greater than 40%, ensuring that only confidently annotated proteins were included. Finally, we removed duplicated sequences from the proteome FASTA files using seqkit’s v0.10.0.1 `rmdedub` command (Shen et al. 2024). For more information on the number of filtered proteins, please see suppl. table 3.

### Differential gene expression analysis

We conducted a quality assessment of the expression data (see suppl. fig. 2) and performed differential gene expression analysis using DESeq2 v1.46.0, analysing each species separately (Love et al. 2014). Count tables were loaded, and a DESeqDataSet object was created with a design formula including genotype-treatment interaction. After setting the treatment reference level and prefiltering genes with low counts, we defined contrast matrices for comparisons such as infected vs. mock across all genotypes, within susceptible or resistant genotypes, and between resistant vs. susceptible genotypes under infected conditions. The `lfcShrink` function was applied to stabilise log fold changes, and DEGs were identified based on an absolute log2 fold change >1 and adjusted p-value ≤0.05.

### Phylogenomics and Orthology

We performed a BUSCO analysis v5.7.0 on the curated proteomes (custom options: `-m protein -l solanales_odb10`) for phylogenetic tree construction (Manni et al. 2021). Subsequently, we used the busco_phylogenomics pipeline to construct species phylogenies, which we visualised with Accurate Species Tree EstimatoR (ASTER*, v1.16). We used Orthofinder v2.5.5 to derive insight into orthologue genes among the different species. We built a central Orthofinder project incorporating the curated proteomes of the five core *Solanum* species and six more distantly related pentapetalae plant species using the same proteome versions as in Sucher et al. (2020). We then used the R package UpsetR v1.4.0 to visualise intersections of shared/unique orthogroups across different scales. In cases where multiple DEGs (e.g., isoforms) were detected per species and orthogroup, we selected those with the most significant and strongest differential expression.

### Co-expression networks (WGCNA)

We constructed weighted gene correlation networks with the R-package WGCNA v1.73 (Langfelder and Horvath 2008). We generated multiple independent networks at the single-copy orthogroup and species levels. To generate the OG network, we first selected all genes of all five species, which were assigned to single-copy orthogroups and renamed them with the respective OG-ID. Then, we generated regularised log-transformed (`rlog`) expression values per sample and merged the per OG expression values of all species into one consensus file, which we used as a starting point for the WGCNA. We analysed both species- and OG-level networks as follows.

First, the data set was inspected for potential outliers by hierarchical clustering using the `hclust()` function v3.6.2 (see suppl. fig. 4A). To derive a soft threshold, we used the `pickSoftThreshold()` function. We evaluated the fit of the scale-free topology model and the mean connectivity (e.g., see suppl. fig. 4B). We then used a custom wrapper of the `blockwiseModules()` function to optimise the settings for each network separately (see suppl. tab. 4, and online resources). Additionally, we used the following custom options: maxBlockSize=nrow(datExpr), networkType=”signed hybrid”, TOMType=”signed”, minModuleSize=30, reassignThreshold=0, checkMissingData=F, replaceMissingAdjacencies=T. We assessed the module assignment using plotDendroAndColors() and tested its robustness by preservation testing (see online resources, and suppl. fig. 4C and suppl. fig. 5).

We investigated per-species module-trait relationships for n modules using linear models accounting for the fixed binomial effects of *genotype* (hence, resistance phenotype) and infection. We excluded the random effect *repetition* to reduce the risk of overfitting the model, as the covariable *repetition* variance approached zero.

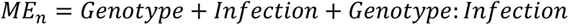

We ranked the plant genotypes according to their LDT values to infer ordinal phenotypic data with the OG network and set the most resistant genotype as a contrast reference.

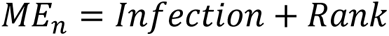

We extracted the model estimates to visualise the OG-module-trait relationships for easier interpretation. Accordingly, we calculated partial- and non-partial η^2^ -values for single-species networks and corrected the p-values with Benjamin-Hochberg correction (Yekutieli and Benjamini 1999).

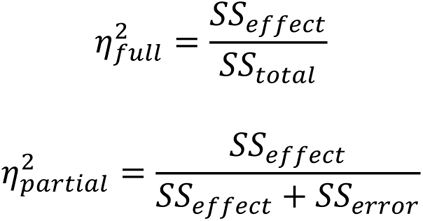

We used a custom script to define hub genes based on eigengene centrality and a dynamic thresholding method which identifies inflexion points of the weight distribution (Tominello-Ramirez et al. 2024). The hub-gene assignment was visually inspected and adjusted if needed (see online resources).

### Gene Regulatory Network Analysis

We conducted a gene regulatory analysis to uncover causal regulatory relationships among the coregulatory genes identified by WGCNA. Accordingly, we discretely predicted transcription factors for all species using the `Transcription Factor Prediction Tool` of the Plant Transcription Factor Database (Jin et al. 2017). Then, we used the R-package GENIE3 v3.20 to construct the Gene Regulatory Network (GRN) using standard settings. We used the same custom script as before to filter low-weight edges and define GRN-hub genes based on eigenvector centrality.

### GO-Term enrichment

To gain insights into the functional framework of the various sets of genes, we performed a gene ontology (GO) analysis on both species-level and cross-species levels. We derived GO terms using PANNZER2 for species-level analysis and converted the results into a BiNGO-compatible format with a custom R-script. We extracted the overlapping genes and GO terms from all species for cross-species analysis. Those were collected in a consensus orthogroup-based BiNGO file. Finally, we performed GO-term enrichment using the Cytoscape v3.10.3 application BiNGO v3.0.5 to interpret the functional implications of defined gene sets.

### Phylostratum analysis

We employed a robust phylostratigraphic approach to classify all transcripts according to their phylogenetic origin, carefully addressing potential biases such as homology detection failure. We used the software GenEra v1.4.2, which uses the DIAMOND algorithm v2.0.14 to align the sequences of protein-coding genes of all five *Solanum* species against the NCBI non-redundant database (as of Sept. 2024). The most distant taxonomic node is considered as the age of this gene family (Barrera-Redondo et al. 2023; Buchfink et al. 2021; Lotharukpong et al. 2024; Schoch et al. 2020; Sayers et al. 2020).

### Transcript Age Index

Following the association of genes with their putative age, we used the R-Package myTAI v0.9.3 to quantify shifts in the transcript age index (TAI) (Drost et al. 2018). *TAI*_*s*_ is defined by the number of treatments (*s*), the relative gene age of *n* genes *i* (*ps*_*i*_) and the expression level of *e*_*is*_, As:

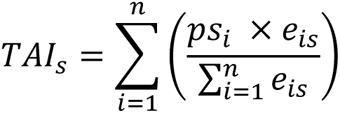

We used normalised log2-transformed read counts (DESeq2 rlog()) as input-expression data.

### Transcript Divergence Index

We employed a dN/dS-based approach to assess evolutionary divergence at the transcript level. After extracting the longest ORFs with TransDecoder v5.7.1, we applied divergence_stratigraphy() from orthologr v0.4.2 (Drost et al. 2015). This method identifies best reciprocal hits via BLASTp, aligns them pairwise with Needleman–Wunsch (Needleman and Wunsch 1970), and converts the protein alignments to codons in PAL2NAL for dN/dS calculations (Suyama et al. 2006; Comeron 1995), using *Solanum melongena* as an outgroup (Wei et al. 2020). The resulting dN/dS values were grouped into deciles (“divergence strata”) to facilitate comparisons with phylostratigraphic data. We used these strata to compute the Transcript Divergence Index (TDI) with myTAI, where *TDI*_*s*_ depends on each gene’s divergence stratum (*ds*_*i*_) and the expression level (*e*_*is*_) as:

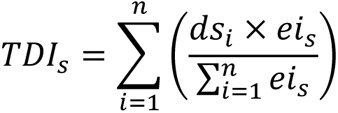

### Statistics and visualisation

We used the programming language R v4.4.0 in RStudio for statistical analysis. In particular, we used packages of the tidyverse v2.0.0, such as tidyr v1.3.1, stringr v1.5.1 and dplyr v1.1.4. All figures were prepared using ggplot2 v3.5.1 and curated for publishing in Inkscape.

### Online resources

Detailed code, including all scripts, documentation and environment information are available at the following GitHub repository: github.com/PHYTOPatCAU/SolanumPhylotranscriptomics

### Data availability

Sequence data and processed read counts from this article are available in the NCBI GEO repository under accession number GSE288242 (see suppl. tab. 14).

## Results

### Wild tomatoes show signs of multiple transcriptome differentiation events

We performed a phylogenomic analysis using curated proteome sequences to corroborate the understanding of the phylogenic relationships between the five tomato species (Solanum - sections Lycopersicon and Lycopersicoides): *S. chilense, S. lycopersicoides, S. habrochaites, S. pennellii,* and *S. pimpinellifolium* (fig. 1A). We found that most protein sequences (13,280 orthogroups of 26,249 total) were shared among all five species, highlighting a strong core proteomic foundation (fig. 1B). In contrast, species-specific unique protein sequences were relatively low, e.g., with *S. pennellii* carrying 2,028 unique proteins and *S. lycopersicoides* 1,795. This pattern underscores the evolutionary conservation within the sections while still allowing room for species-specific adaptations. To further characterise the proteomic landscape’s evolutionary history, we employed phylostratigraphy to map the age of genes across all tested species.

**Figure 1:**
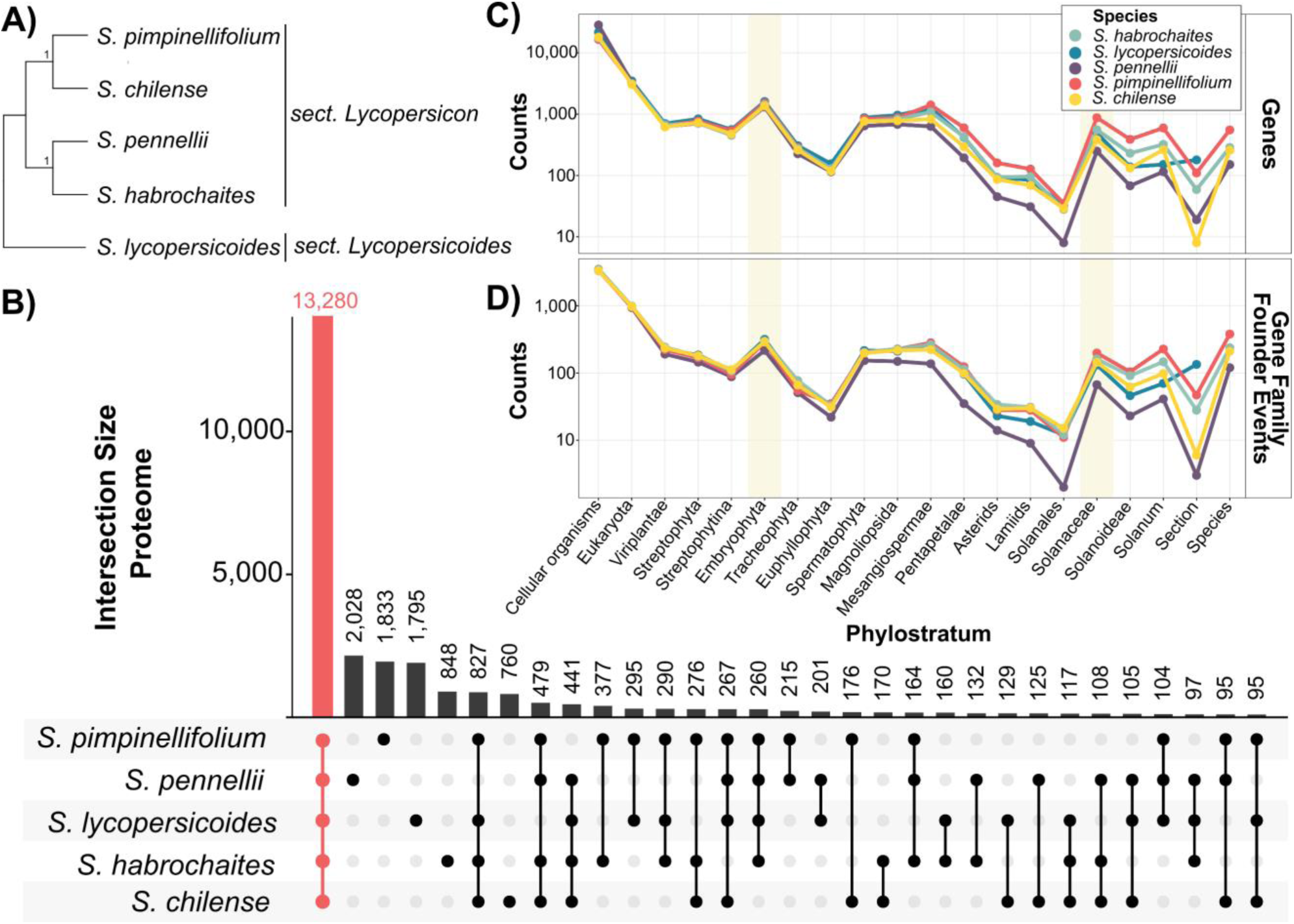
Phylogenomic Analysis Reveals a Highly Conserved Proteome and Recent Differentiation Among Five Solanum Species. **A)** Representative phylogenetic tree of the five tomato species based on BUSCO genes. Numbers represent branch support values based on quadripartition, considering the four clusters surrounding a branch. **B)** Orthofinder-based Upset plot of the proteomes, indicating that a large fraction of proteins is shared among species (in red), with only a small subset unique to individual species. Phylostratigraphic maps display the number of genes **(C)** and gene family founder events **(D)** across the tree of life. Shaded regions highlight phylostrata corresponding to the peaks of genetic innovation during the development of land plants and within the Solanaceae. Counts correspond to the number of genes or gene family founder events, respectively.

By constructing phylostratigraphic maps, we can visualise the age of the genes and when key gene families emerged. This allows for assessing whether genes are conserved or have diversified across lineages. As anticipated, all five species exhibit a strongly conserved progression in the number of genes with a certain age (fig. 1C) and the number of associated gene family founder events (fig. 1D). Most genes trace their origins back to the emergence of cellular organisms (e.g., 18,056 *S. chilense* genes from 3,346 gene family founder events [GFFE]), with a marked decline in new gene origins over time. However, notable spikes occur during key evolutionary milestones: the development of land plants (Embryophyta, 290 GFFE, 1,362 new genes in *S. chilense*) and the rise of flowering plants (from Spermatophyta to Mesangiospermae, in sum 2,371 new genes and 637 GFFE in *S. chilense*). Following these periods, gene assignments gradually decrease until the clade Solanaceae, where there is a renewed burst of new gene families (*S. chilense*: 144 GFFEs). All species harbour a modest number of phylostratigraphically young genes (*S. chilense*: 210 GFFE), which may indicate ongoing species-specific differentiation or bursts of gene family innovation since the emergence of Solanaceae. However, these putatively young genes must be critically assessed, as they may partly reflect technical artefacts (such as reference genome annotation quality or analytical issues like homology detection failure).

The high number of shared orthogroups and remarkably similar age estimates for gene family founder events until the clade of Solanum illustrate the strongly conserved evolutionary history of the five wild tomato species. This conservation suggests that, despite species-specific innovations, their core genomic and evolutionary trajectories have remained remarkably stable.

### *Sclerotinia sclerotiorum* tolerance is highly diverse among and within wild tomato species

We previously performed a study to identify genotypes differing in disease resistance against the fungal generalist-pathogen *S. sclerotiorum* (Einspanier et al. 2024). Generally, we observed a wide diversity in QDR phenotypes with strong species- and accession-specific patterns.

Accordingly, we observed that the lesion growth rate (denominated Lesion Doubling Time, LDT) was generally shorter on *S. pennellii* and *S. pimpinellifolium* genotypes (i.e., pathogen growth is faster). At the same time, *S. lycopersicoides* accessions appeared relatively resistant (fig. 2A). We found the biggest quantitative differences between genotypes of the same species for *S. lycopersicoides* (lsmean_susceptible_ = 6.80 h vs. lsmean_resistant_= 7.88) and *S. pennellii* (lsmean_susceptible_ = 5.86 h vs. lsmean_resistant_ = 6.65h, see suppl. tab. 1).

**Figure 2:**
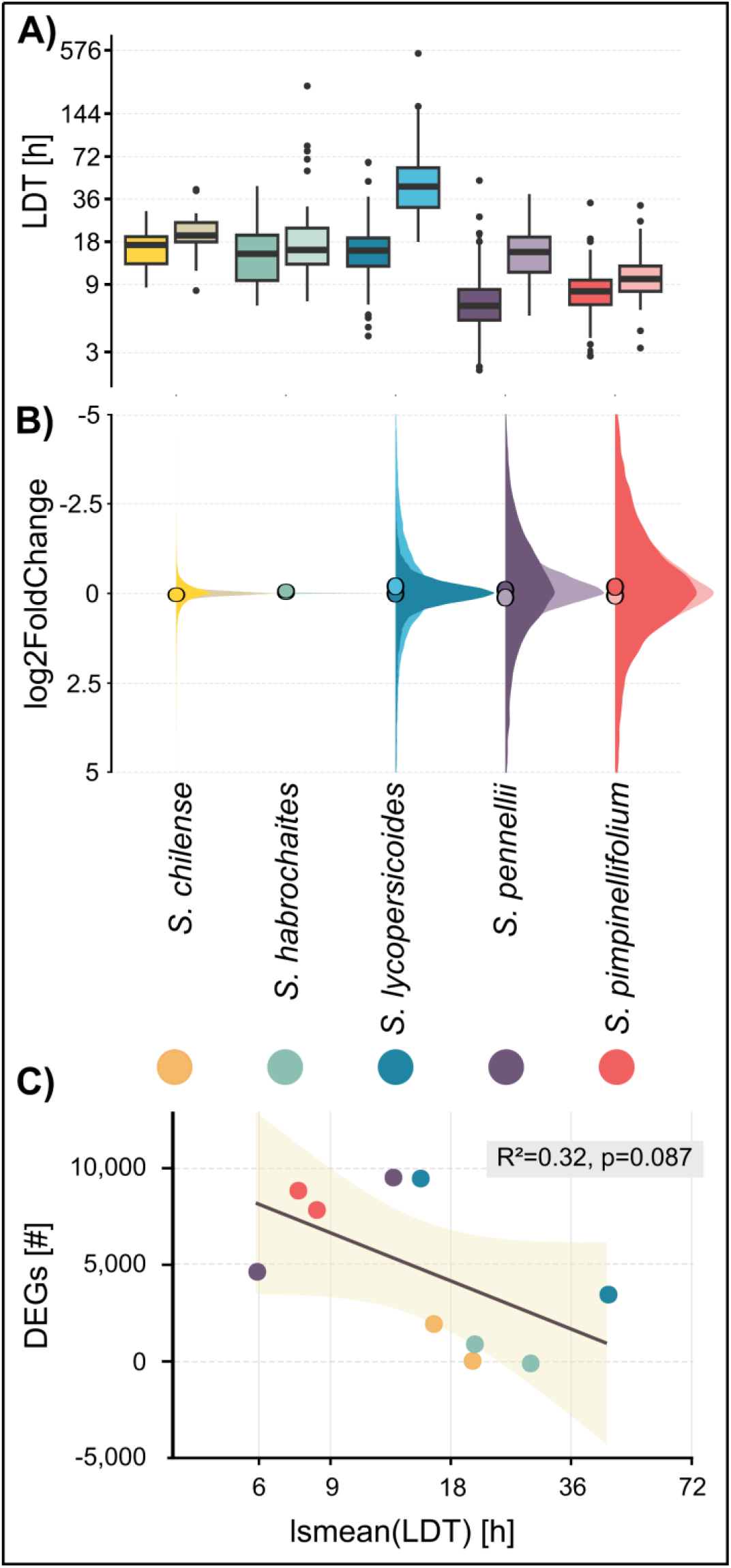
The inoculation with *S. sclerotiorum* leads to different levels of susceptibility and heterogeneous transcriptomic responses. **A)** Lesion Doubling Time (LDT) values in hours for five host species (two genotypes each) serve as a quantitative measure of the level of quantitative disease resistance (QDR). Data adapted from Einspanier et al. (2024). **B)** Density plot of log₂FoldChange in gene expression across all accessions following infection with *S. sclerotiorum* compared to mock treatment, illustrating the overall transcriptomic response to infection. Dots represent the mean log₂FoldChange. **C)** Pearson correlation analysis testing the interaction between the number of differentially expressed genes by genotype (induced by infection) and levels of resistance (LDT). The shadow represents the SD.

### *S. sclerotiorum* infection-induced shift in gene expression underlies genotype specificity

We then conducted gene expression profiling on detached leaves during the lesion growth phase, contrasting accessions with different LDT. An average of 4.3 mio. reads mapped to the respective reference genomes, with a mean mapping rate between 74% and 89% (mapping statistics in suppl. tab. 2). We performed differential gene expression analysis by genotype comparing infected versus control conditions. Interestingly, we observed strong variation in differential (infection-induced) gene expression levels between the species. We observed few weakly differentially expressed genes for both *S. chilense* (sus. genotype: 2,104 DEGs, res. genotype 194 DEGs, fig. 2B). and *S. habrochaites* genotypes (sus. genotype: 1,058 DEGs, res. genotype 65 DEGs) and a stronger regulatory response by the other host species (*S. pennellii*: sus. genotype: 4,808 DEGs, res. genotype 9,685 DEGs; *S. lycopersicoides*: sus. genotype: 3,627 DEGs, res. genotype 9,635 DEGs; and *S. pimpinellifolium*: sus. genotype: 9,007 DEGs, res. genotype 8,007 DEGs, see suppl. tab. 5). We performed a Pearson correlation analysis to test the relationship between the level of QDR and the absolute number of differentially expressed genes and observed a non-significant, negative association between the two parameters (p=0.087, R²=0.32, fig. 2C). However, nested species effects with contrasting trends might not be reflected sufficiently by this analysis, as we measured a lower number of DEGs on the resistant *S. lycopersicoides* genotype vs. the susceptible genotype and the opposite trend for *S. pennellii.* These findings suggest that the absolute number of genes reprogrammed during infection alone does not explain QDR variability at this level. We also found no relationship between sequencing/mapping depth and the number of DEGs (see suppl. fig. 1).

### Genotypes of the same species show strong regulatory plasticity

Seeing the results above and knowing that constitutively expressed defence-associated genes might also govern QDR, we performed a differential gene expression analysis contrasting the resistant vs. the susceptible genotype by species in infected conditions. This allows the identification of infection-induced regulatory features and the expression of non-induced genes alike. Interestingly, the number of differentially expressed genes between the two genotypes varies strongly between the species (1,438 DEGs up and 1,644 DEGs down in *S. habrochaites* versus 5,855 DEGs up and 4,962 DEGs down in *S. lycopersicoides*). While we found strong alterations in the magnitude of differential gene expression between two *S. lycopersicoides* genotypes (5,855 sign. up-regulated genes in the resistant genotype, 4,962 downregulated, respectively), *S. pimpinellifolium* genotypes differed on a much smaller scale (1,483 up-regulated DEGs, 1,644 downregulated DEGs). Intermediate levels of differential expression between *S. pennellii* genotypes, in contrast to strong differences in *S. lycopersicoides,* clearly illustrate that the order of magnitude in gene expression does not explain the level of resistance or susceptibility (fig. 3A).

**Figure 3:**
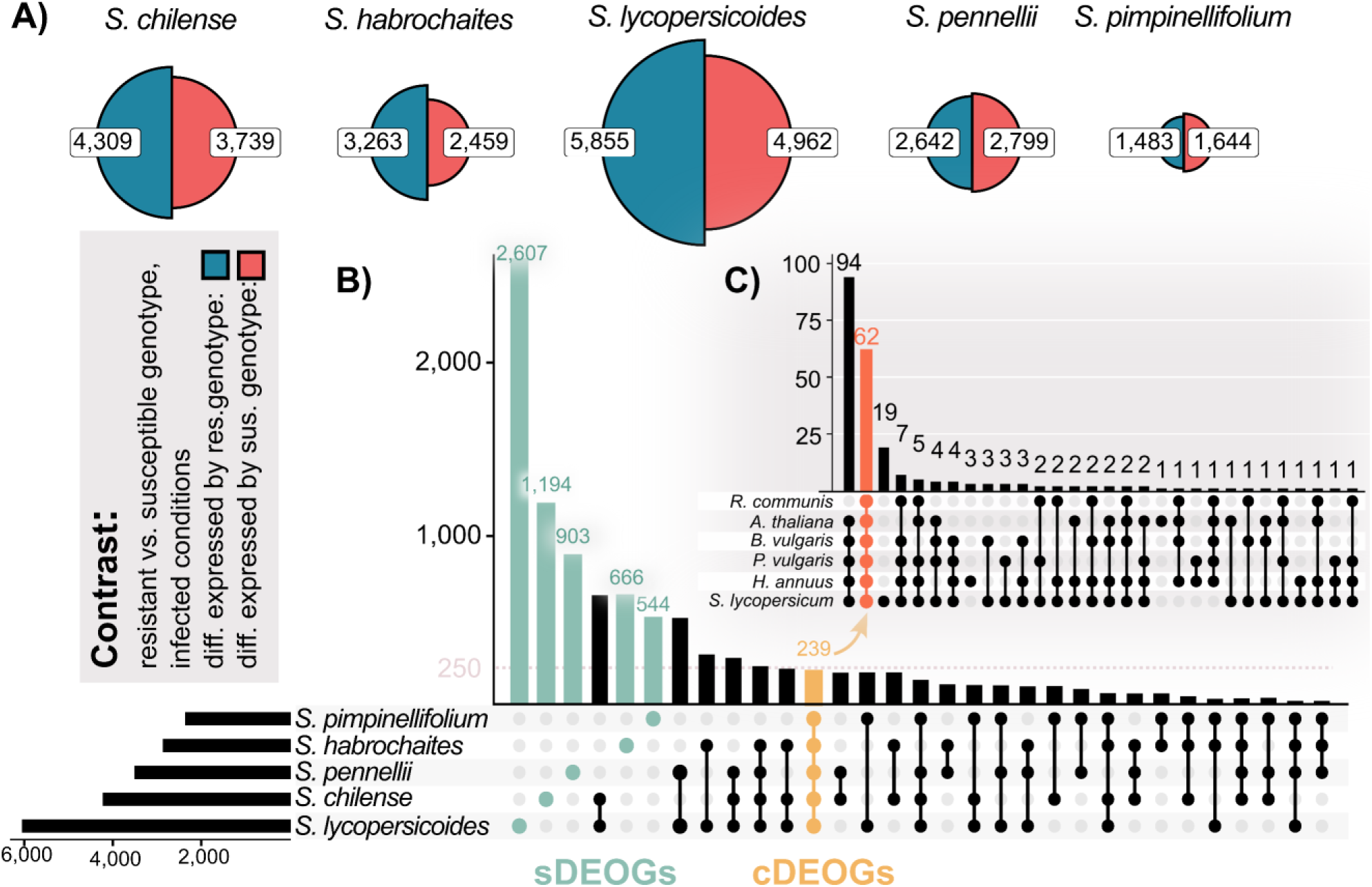
Species-Specific Differential Gene Expression Related to Varying QDR Levels. **A)** Pie charts showing the number of genes differentially regulated between genotypes with varying levels of quantitative disease resistance (QDR) within each species. DEGs were defined between genotypes with differing levels of QDR in infected conditions. **B)** Upset plot illustrating the overlap of between-genotype differentially expressed orthogroups across the five tomato species. This intersection distinguishes species-specific differentially expressed orthogroups (sDEOGs) from a set of shared core differentially expressed orthogroups (cDEOGs). **C)** Upset plot presenting the number of differentially expressed core DEOGs (cDEOGs) observed in other Pentapetalae species upon *S. sclerotiorum* infection, highlighting the conservation and variability of these core genes across a broader phylogenetic context. Data adapted from Sucher et al. (2020).

Next, we tested whether resistance- or susceptibility-DEGs are unique or shared among species. For this, we selected all DEGs between resistant and susceptible accessions within a species and conducted a membership analysis based on gene orthologs to classify differentially expressed orthologs (DEOGs). We observed that most DEOGs were unique to the respective species. Accordingly, we measured 2,607 specific DEOGs (sDEOGs) for *S. lycopersicoides,* 1,194 sDEOGs for *S. chilense* and 903 sDEOGs from *S. pennellii.* However, we also identified 239 core DEOGs (cDEOGs) shared among all five tomato species, indicating that those genes are differentially expressed between the two genotypes of all species. Interestingly, we could show that the majority of those cDEOGs is differentially expressed upon *S. sclerotiorum* inoculation in six further related pentapetalae plants: 62 of the cDEOGs are induced in all tested plant species (*Ricinus communis*, *Arabidopsis thaliana, Beta vulgaris, Phaseolus vulgaris, Helianthus annuus* and *S. lycopersicum*), and 94 cDEOGs are infection-induced in all plants of this set excluding *R. communis.* We further identified 19 cDEOGs, which might be specific to the genus Solanum (fig. 3C). These findings illustrate that only a small proportion of all DEOGs show a conserved expression pattern over pentapetalae plants. At the same time, regulatory differentiation between genotypes with contrasting QDR appears to be mostly specific to the nested species.

In the pentapetalae plant set, most conserved differentially expressed orthologous genes (cDEOGs) are significantly induced by infection. However, within our wild tomato dataset, only a small fraction of these genes (n=16) is both infection-induced and vary significantly between genotypes of differing QDR. Notably, most cDEOGs exhibited pronounced presence-absence patterns in their infection response (see suppl. fig. 3), suggesting that their regulation is highly variable across species. Species-specific DEOGs also display differing regulatory profiles: While ∼60% of *S. pimpinellifolium*, *S. pennellii*, and *S. lycopersicoides* sDEOGs are induced by infection, only a marginal fraction of sDEOGs in *S. chilense* or *S. habrochaites* is induced (suppl. fig. 3). This pattern underscores the regulatory flexibility and diversity among wild tomato species in their response to pathogenic infection, likely reflecting a wide range of adaptive strategies in their defence mechanisms.

### The regulation of induced cDEOGs is diversifying among Solanum species

We characterised the function of the 16 infection-induced cDEOGs (see suppl. tab. 6). Interestingly, we identified two genes (OG0006904 and OG0018399) matching transcription factors of the AP2/ERF family. One of them, OG0018399, was previously characterised as an *S. lycopersicum* ERF-D6 transcription factor (Tominello-Ramirez et al. 2024). Furthermore, we found a putative chitinase/cell-wall degrading enzyme (OG0005084) and a putative UDP-glycosyltransferase (OG0000514) among oxidative stress regulation (fig. 4).

**Figure 4:**
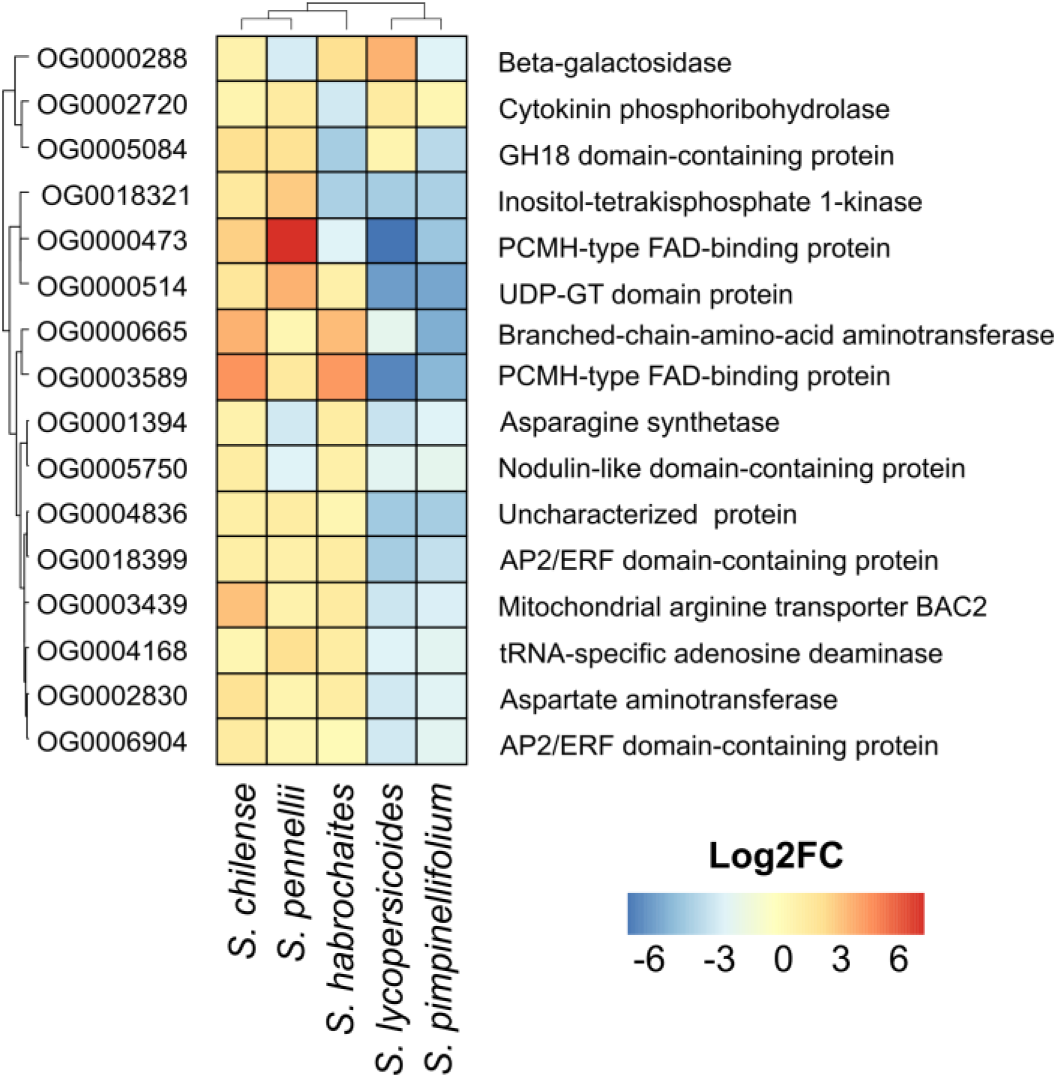
Differential Expression of 16 core infection-induced DEOGs. The heatmap illustrates the differential expression levels of 16 core infection-induced differentially expressed orthologous genes (DEOGs) across the five tomato species. Each row represents a gene, and columns correspond to comparisons between resistant and susceptible genotypes within each species. The colors indicate the magnitude and direction of expression differences in infected conditions, showing inconsistent gene regulation patterns associated with resistance and susceptibility to infection.

We hypothesised that the expression pattern of those infection-induced cDEOGs might be mostly uniform, as they are induced by infection and differentially regulated between susceptible and resistant genotypes. However, we observed a diversified regulatory pattern among all induced cDEOGs with contradictory regulation. Most interesting, the putative ERF D6 (OG0018399) is significantly upregulated in the resistant genotypes of *S. habrochaites, S. pennellii* and *S. chilense*. Yet, it is downregulated in the resistant genotypes of *S. pimpinellifolium* and *S. lycopersicoides* (indicating a higher induction in the susceptible genotypes, fig. 4). Interestingly, except for OG0004836, all induced cDEOGs share an evolutionary origin predating the emergence of flowering plants, suggesting that these highly conserved genes remain subject to differential regulation (suppl. tab. 6). Concluding, our differential gene expression (DEG) analysis gained initial insights into the regulatory dynamics driven by contrasting infection conditions and/or genotypes and shows that rewiring of expression pattern of shared infection-induced DEOGs may be source of QDR adaptation at the species level.

### Weighted Gene Correlation Network Analysis reveals evidence of transcriptome specification

While informative, DEG analysis alone cannot fully capture the complexity of the underlying regulatory pathways and networks. Thus, to further investigate global transcriptional rewiring of QDR responses in our five species and to achieve a more comprehensive understanding of QDR across species, we employed multiple Weighted Gene Correlation Network Analyses (WGCNA).

First, we constructed an orthology-based pan-species network. We hypothesised that the overarching gain of QDR might be correlated with the module eigengenes. To test this, we constructed a gene-correlation network based on 7,419 single-copy orthogroups, which were grouped into eleven distinct coregulatory modules, with a total of 462 hub genes (fig. 5A&B, suppl. fig. 4). Following network construction, we examined both inter- and intramodular edges. We validated the biological relevance of module assignments through functional assignments using Gene Ontology terms. We used a linear model to infer QDR-phenotypes as ranks with the module eigengenes (MEs).

**Figure 5:**
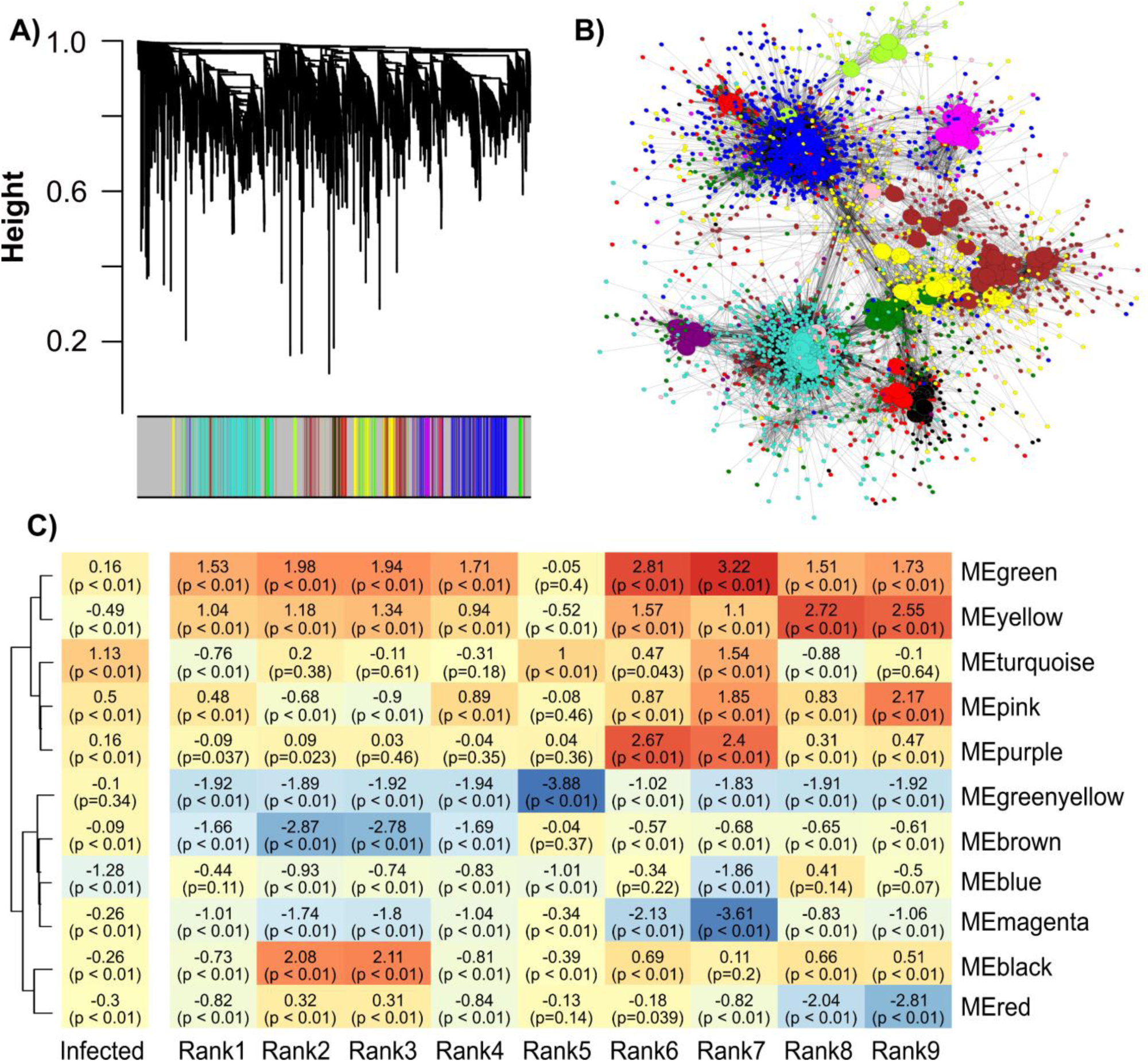
Weighted Gene Correlation Analysis of a single-copy orthogroup across five tomato species. **A)** Dendrogram illustrating the clustering of single-copy orthogroups into modules, with each module assigned a unique color. **B)** Network visualization of the orthogroup-based modules, where large nodes represent highly connected hub genes. **C)** Module-trait relationships derived from a linear model using module eigengenes as the response and infection state plus resistance rank as predictors. Resistance rank is relative to the most resistant genotype (*S. lycopersicoides*, LA2951), and infection status is compared to mock conditions (see suppl. tab. 1). The coefficients shown represent effect size estimates from ANOVA, while the accompanying p-values indicate significance after FDR correction.

Interestingly, we found no significant and biologically meaningful association of module eigengene expression with the LDT. While certain modules (e.g., blue, yellow, turquoise) showed a strong association with the infection as such, there was no clear gradient in module eigengene estimates from the most susceptible to the most resistant genotype. Moreover, despite having similar LDT (Rank7/Rank8, suppl. tab 1), these two genotypes show notably contrasting module eigengenes in infection-associated modules (for instance, Rank7 blue = -1.86 versus Rank8 blue = 0.41). At the same time, the strongest associations in the module-trait relationships emerge between genotypes within the same species (such as Rank2 and Rank3 or Rank1 and Rank4), suggesting that QDR regulation could be tightly integrated into genotype- or species-specific regulatory frameworks.

Accordingly, we hypothesise that the OG-based network (based on two genotypes per species) may lack the statistical power to link regulatory patterns to cross-species phenotypes confidently or that phenotypic resistance is achieved through interlinked, species-specific pathways (fig. 5C, suppl. tab. 7).

### Species-specific network topology is linked to QDR variation

We, therefore, performed per-species regulatory network analyses to gain higher-resolution insights into QDR regulation. Specifically, we compared *S. lycopersicoides* and *S. pennellii* - the two species that displayed the most pronounced differences in resistance phenotypes across the tested genotypes. In *S. pennellii*, 16,577 genes were clustered into eight co-expression modules, while 18,558 *S. lycopersicoides* genes were grouped into nine modules (fig. 6A&B). For both species, we identified modules with predominant genotype association (exhibiting no significant interaction with infection), thus reflecting basal, genotype-specific alteration of gene expression (e.g., black and green modules in *S. pennellii* and the brown and yellow modules in *S. lycopersicoides,* fig. 6C&D). Conversely, we detected six *S. pennellii* and seven *S. lycopersicoides* modules significantly associated with infection (estimate > 0.2). The modules showing a significant association with the infection:genotype interaction are particularly interesting, as these are likely the strongest candidates for explaining the differential QDR phenotypes. Complementary to the statistical analysis, we evaluated the module eigengenes (MEs) visually and used functional annotation to identify modules with a putative role in resistance (suppl. fig. 6&7, suppl. tab. 8&9). Accordingly, we identified three modules in *S. pennellii (*‘red,’ ‘pink,’ and ‘blue’) and three *S. lycopersicoides* modules (‘pink,’ ‘green,’ and ‘turquoise’) with a putative role in resistance. Next, we combined both datasets using orthogroups to quantify the extent of overlap between resistance modules of the respective species. We assessed the robustness of this overlap using Fisher’s exact test with FDR-corrected p-values. 2,264 of 3,683 orthologous *S. pennellii* resistance genes were significantly enriched in the *S. lycopersicoides* resistance modules (suppl. tab. 10&11, fig. 6E&F). However, a significant number of *S. pennellii* genes did not cluster in *S. lycopersicoides* resistance modules: 386 genes were assigned to the yellow module and 240 to the green module. Interestingly, 1,277 genes from the *S. pennellii* resistance modules could not be assigned to a specific *S. lycopersiocides* module and were assigned to the grey module. We observed a similar pattern when projecting genes from the *S. lycopersicoides* resistance modules onto the *S. pennellii* gene regulatory network. In this case, 1,918 of the 3,702 orthologous genes were located in the *S. pennellii* resistance-associated blue module. In comparison, a significant number of genes (1,101 genes) was assigned to the brown module, which is specific to infection response rather than resistance. These findings indicate that although several QDR-related genes might be shared across species, a significant subset of QDR genes remains associated with distinct non-resistance modules reflecting species-specific regulatory fine-tuning.

**Figure 6:**
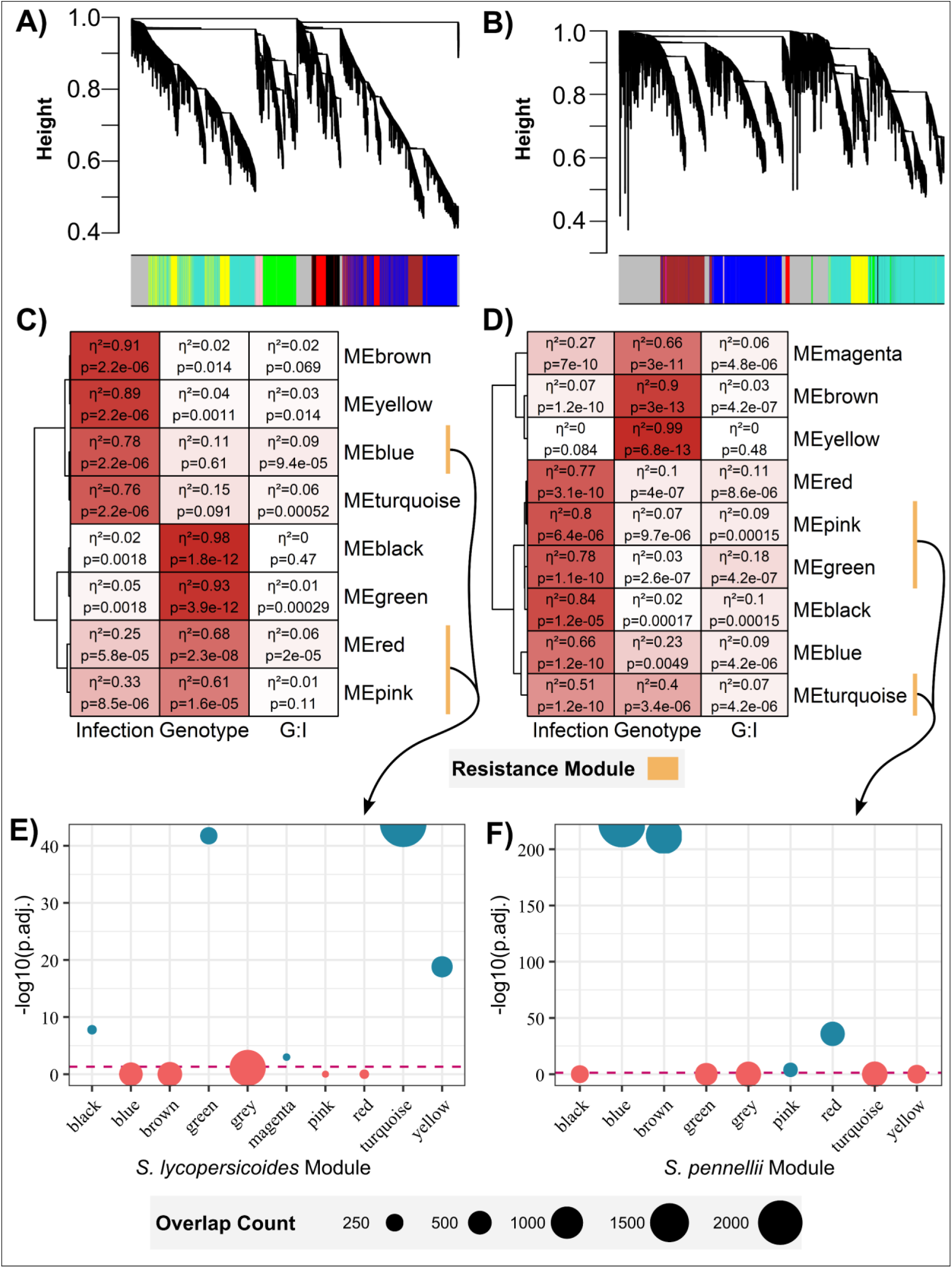
Weighted Gene Correlation Network Analysis of A) *S. pennellii* and B) *S. lycopersicoides* RNAseq data. Module eigengenes were correlated with the traits “Infection”, “Genotype”, and the interaction “Genotype x Infection” using a linear model. The effect size is provided as η^2^, and FDR-corrected p-values are provided for **C)** *S. pennellii* and **D)** *S. lycopersicoides*. Resistance-associated modules were marked. The module preservation of **E)** *S. pennellii* resistance genes in *S. lycopersicoides* modules and **F)** *S. lycopersicoides* resistance genes in *S. pennellii* modules were tested using orthogroup-based inference and a Fisher’s exact test. The dot size indicates the number of overlapping genes, while the blue colour indicates a significant enrichment.

### Functional enrichment reveals a broad core functional set and specific species functions

We then performed a GO-term enrichment analysis on both species’ shared and unique QDR-module gene sets to assess whether those harboured specific functional traits. Among the 2,264 overlapping genes, we identified a diverse array of significantly enriched GO-terms clustered into six groups: primary/secondary metabolism, localisation, developmental processes, protein metabolism, signalling, and response to stimulus. Within these clusters, we found enrichment of protein transport and localisation, lipid and carbohydrate metabolism, and cinnamic acid synthesis involving processes. Notably, we did not observe a clear enrichment for hormone homeostasis (fig. 7A). In contrast, the *S. pennellii*-specific genes were clustered into four functional groups: biosynthesis (including macromolecule/protein glycosylation), positive regulation of cellular/biological processes, stress response, and catabolism, with modest enrichment for signalling functions (fig. 7B).

**Figure 7:**
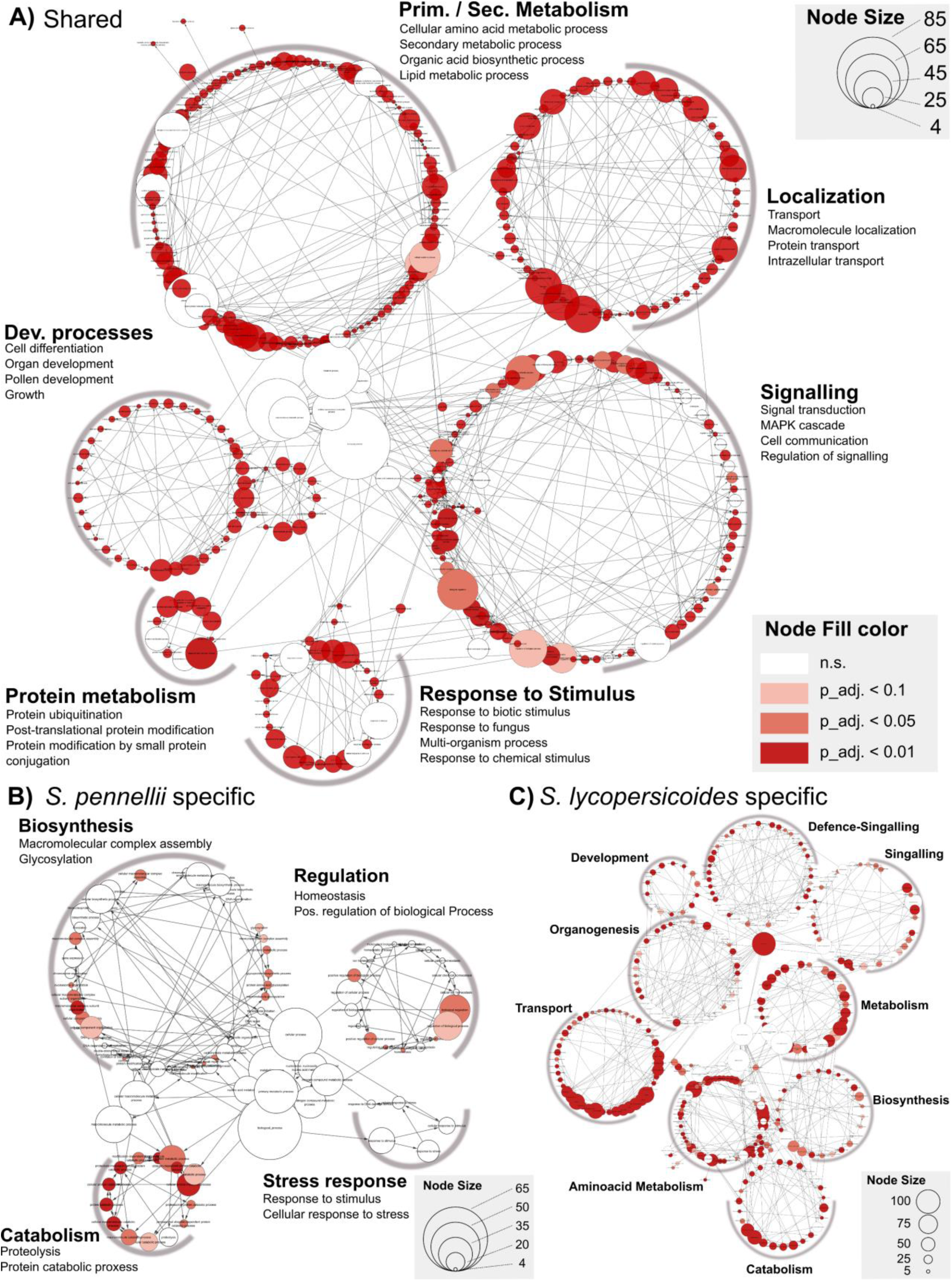
Gene Ontology (GO) Enrichment Analysis of Resistance Coexpression Modules. **A)** Core genes shared across the resistance modules of *S. pennellii* and *S. lycopersicoides.* **B)** *S. pennellii* -specific genes, **C)** *S. lycopersicoides-*specific genes. In each panel, the size of a dot corresponds to the number of genes associated with a GO term, while the fill colour indicates the FDR-corrected significance level of the enrichment. This visualisation highlights key functional categories enriched in core and species-specific resistance responses.

For *S. lycopersicoides,* we identified nine differentially regulated functional clusters, with a particular emphasis on transport, amino acid metabolism, general metabolism, and defence signalling (fig. 7C). These findings support a two-tier model of QDR regulation between species: a broad, shared set of genes provides general disease resistance, while host-specific mechanisms offer more targeted functions. For example, *S. pennellii* may rely on detoxification via glycosylation, whereas *S. lycopersicoides* could depend more on specialised transport processes. This model underscores how an equilibrium of shared and species-specific strategies may shape effective defence responses across different *Solanum* species.

### Shared QDR regulation comprises evolutionarily conserved genes

To test the hypothesis that the specification of QDR signalling may be subject to diversifying selection, we calculated the Transcript Age Index (TAI) and Transcript Divergence Index (TDI) for both shared and species-specific gene sets. Our analysis revealed that species-specific genes are evolutionarily younger, as suggested by increased TAI (e.g., *S. pennellii*: overlap TAI = 1.69 vs. unique TAI = 1.92; *S. lycopersicoides*: overlap TAI = 1.73 vs. unique TAI = 2.4, see table 1). Additionally, shared genes from resistance modules showed lower TDI values, indicating signs of stronger purifying selection than observed on the unique sub-modules (*S. pennellii*: 4.3 vs 4.7; *S. lycopersicoides:* 4.3 vs 4.9, see suppl. tab. 12). This is further supported by significantly elevated ratio of nonsynonymous mutations to synonymous mutations (dNdS) in each species’ uniquely regulated gene set compared to the whole resistance modules (suppl. fig. 8). Accordingly, we hypothesise that both species share a conserved, ancestral suite of genes that presumably form the “backbone” of the quantitative defence network regulation, while younger, less strongly conserved genes underlie specification. This balance between conservation and innovation might be critical for the dynamic fine-tuning of defence responses against generalist pathogens over evolutionary timescales.

**Table 1:** Per-Species Transcript Age Index (TAI) of Resistance-Modules covering shared and unique gene sets. Mean TAI values are provided with +/- SD.

| Species | Whole Resistance Modules | Overlap | Unique |
| --- | --- | --- | --- |
| *S. pennellii* | 1.77319975 ± 0.006335785 | 1.6909445 ± 0.00817666 | 1.922385 ± 0.007211685 |
| *S. lycopersicoides* | 1.981305 ± 0.019387067 | 1.7368255 ± 0.009673017 | 2.4065145 ± 0.035387022 |

### QDR in *S. pennellii* genotype LA2963 may have evolved by co-optation of a transcription factor

To test the above hypothesis whether regulatory hubs or the periphery drive the evolutionary signature of the network, we integrated gene-regulatory network analysis with dN/dS analysis across multiple species. We extracted potentially interesting transcription factors according to the following criteria: a) hub in the WGCNA network (high connectivity), b) gene in the resistance module, c) hub in the gene regulatory network and identified highly connected transcription factors for both focal species (18 in *S. lycopersicoides* and 10 in *S. pennellii)*. We then focussed on TFs exhibiting differential expression patterns in response to infection and between genotypes with varying resistance levels and compared their subnetworks across multiple species.

In *S. pennellii,* this subset includes a putative Trihelix transcription factor GT-3b (Sopen09g001470, OG0007365), a MADS-box domain containing-protein (Sopen10g006210, OG0015518) and NAC transcription factor 29 (Sopen05g003630, OG0005445, fig. 8A). In *S. lycopersicoides*, we observed an ethylene-response transcription factor 14 (Solyd03g050610, OG0003738), and two HSF-type DNA-binding domain-containing proteins (Solyd02g064330, OG0000116, Solyd09g071070, OG0009560) with a strong association with the resistant genotype (fig. 8C). Generally, these TFs were more strongly induced in the resistant genotype and are part of a resistance-associated coregulatory module.

**Figure 8:**
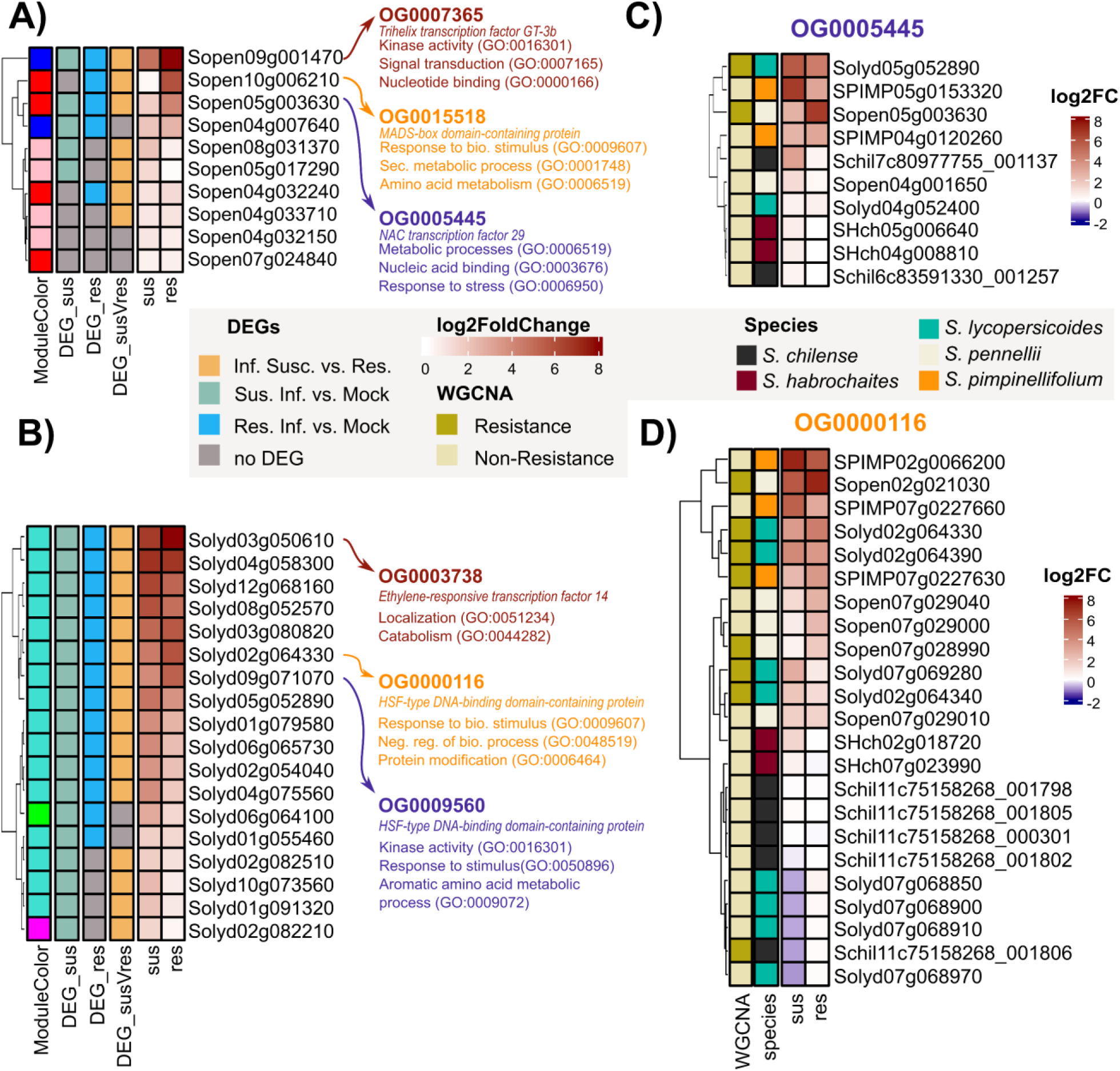
Gene Regulatory Network Analysis Reveals a Shared Transcription Factor Co-opted for Divergent Regulation and Coexpression. The heatmaps in panels **A)** and **B)** illustrate transcription factor (TF) regulation in susceptible and resistant genotypes of *S. pennellii* and *S. lycopersicoides*, respectively. Each TF is a hub gene in the GRN and WGCNA and is located in a resistance module. Each heatmap row corresponds to a TF, with the ‘ModuleColor’ column indicating its assignment to a specific WGCNA resistance module. Differential expression analysis of relevant contrasts is also displayed, with the three most strongly regulated TFs highlighted along with their corresponding orthogroups. For these key TFs, summarized GO terms for their downstream coexpressed genes are provided, offering insights into their functional roles in resistance pathways. The regulation of these resistance-associated TFs (**(C)** *S. pennellii:* Sopen05g003630-OG0005445, **(D)** *S. lycopersicoides: Solyd02g064330-OG0000116*) was then compared across species using orthogroups. The expression of each gene within the respective orthogroup was visualised across varying levels of QDR (susceptible | resistant), with colour coding indicating the respective species and the assignment of each TF to its species-specific resistance module.

We then examined whether the orthologues of these putative resistance-related transcription factors (TFs) showed similar patterns in the other species. We discovered a wide diversity in TF expression among all members of the same orthogroup. For example, the TF Sopen05g003630 (OG0005445) was strongly induced only in the resistant genotype of *S. pennellii*, although this gene is present and expressed in other species. Moreover, OG0005445 is not specifically induced in resistant genotypes nor assigned to a resistance module of the other species. This suggests that the regulation of Sopen05g003630 (OG0005445) might represent a resistance mechanism unique to the *S. pennellii* – *S. sclerotiorum* interaction. So, the regulation and the downstream processes of this TF might have shifted in *S. pennellii* compared to the other species (fig. 8A&B). The genes Sopen10g006210 (OG0015518) and Sopen09g001470 (OG0007365) show an elevated expression pattern specific to the resistant *S. pennellii* genotype. While we could not detect a strong expression of Sopen10g006210 in the other species, the expression pattern of Sopen09g001470 is not linked to the QDR phenotypes (suppl. fig. 9). We observed a similar pattern of diversifying regulation on the *S. lycopersicoides* genes Solyd03g05610 (OG0003738) and Solyd09g071070 (OG0009560), with no clear interspecific QDR-regulation (suppl. fig. 9).

In contrast, the *S. lycopersicoides* gene Solyd02g064330 (OG0000116) appeared highly conserved across species with high similarity in expression pattern among the other species (orthologues also belong to resistance modules and are more highly expressed in resistant genotypes). Even so, we found strong differential regulation depending on the individual paralog (fig. 8D). Interestingly, low dN/dS ratios suggest that both, Sopen05g003630 and Solyd02g064330, transcription factors are under purifying selection (Solyd02g064330 dN/dS = 0.4539 and Sopen05g003630 dN/dS = 0.1884) (suppl. tab. 13). Additionally, the GRNs of all resistance TFs are enriched for younger genes that emerged with the flowering plants and show a significant under-representation of older genes compared to the entire transcriptome (suppl. fig. 10). These younger genes and the low dN/dS ratios do not fully explain the novel regulatory functions. Therefore, we propose that regulatory evolution, rather than changes in nucleotide sequences, drives the evolution and short-term adaptation of these TFs.

## Discussion

In this study, we asked whether variance in observed quantitative disease resistance phenotypes against generalist pathogens in closely related yet diverse species is governed by one or multiple mechanisms and how such variance can evolve. Singular studies on gene or gene family evolution alone cannot fully capture this complexity (Kahlon and Stam 2021). Therefore, we performed integrated transcriptomics across five wild tomato species infected with *S. sclerotiorum*. By combining differential expression, WGCNA, GRN, and deeper evolutionary analysis, we uncover both conserved and species-specific QDR strategies, highlighting a complex interplay between core and adaptive regulatory mechanisms.

### Resistance against *S. sclerotiorum* underlies a complex regulatory interplay

Studies on diverse crops and model plants (e.g., soybean, rapeseed, sunflower, bean, pepper, tomato, *A. thaliana*) reveal broad variation in QDR against *S. sclerotiorum* (Boland and Hall 1994; Boudhrioua et al. 2020; Chauhan et al. 2020; Ding et al. 2021; Fusari et al. 2012; Mazumdar 2021; Mei et al. 2012; Musa-Khalifani et al. 2021; Sucher et al. 2020; Uloth et al. 2013; Yanar and Miller 2003; Einspanier et al. 2024). Accordingly, QDR may be governed by a complex array of regulatory and genomic features, as suggested by the large variation in differentially expressed genes (DEGs), ranging from a few dozen to over 20,000, observed under diverse experimental conditions (Fass et al. 2020; Joshi et al. 2016; Tang et al. 2023; Wu et al. 2016; Sucher et al. 2020; Chen et al. 2022). However, our findings highlight that the number of DEGs following *S. sclerotiorum* infection is not directly linked to QDR levels within or between species. In a broad-scale study, Sucher et al. (2020) proposed that regulatory flexibility and specificity of gene expression changes are essential for QDR. We show that the differential expression of resistance-associated genes is highly diverging between the species, and shared genes show contrasting regulatory patterns among the species. The wide variability of DEG counts and -regulation, as well as the dominance of species-specific patterns, underscores the complexity of plant-pathogen interactions and suggests the role of distinct molecular strategies.

### QDR-Regulation is based on both host-specific and conserved genes

We examined both interspecific and intraspecific regulatory responses to infection. Comparing gene regulatory and co-expression networks across species remains challenging, and cross-species mapping doesn’t allow the identification of species-specific genes. Therefore, we directly compared differentially expressed genes (DEGs) and network topologies by focusing on orthologous genes (Külahoglu et al. 2014; Langfelder et al. 2011; Ovens et al. 2021; Hu et al. 2016). This approach preserves a larger number of species-specific transcripts and facilitates the projection of findings onto more distantly related plant species. Furthermore, leveraging orthologs maintains the integrity of each species’ unique transcriptome and enhances our ability to uncover both conserved and unique regulatory mechanisms that underpin similar phenotypic outcomes.

Although the tested genotypes share notable genome/proteome similarity, the regulatory networks remain highly species-specific, suggesting that fine-scale shifts in gene regulation may drive the emergence of unique traits (Halfon 2017; Smith et al. 2014). In some cases, genotype effects surpass those induced by infection, suggesting genotype-specific signalling dominates in certain backgrounds. This aligns with our WGCNA-based observation that regulatory patterns correlate more strongly with species identity than any uniform resistance gradient. Nevertheless, each species possesses co-regulatory modules linked to resistance. Focusing on the overlap between *S. pennellii* and *S. lycopersicoides*, we found that shared modules appear involved in common defence responses. In contrast, unique sub-modules focus on specialised traits (e.g., signalling or detoxification). This suggests that GRNs are finetuned on a species level, also when it comes to defence against broad-host-range necrotrophic pathogens (Smith et al. 2014). Other indications of GRN specialisation for supposedly generic tasks exist: only a minority of *S. lycopersicum* cis-regulatory elements controlling wound response overlap with *S. pennellii* (Liu et al. 2018; Jones and Vandepoele 2020).

### Rewiring Gene Networks as an Adaptive Strategy

The pattern of gene expression can change faster than the divergence of nucleotides. This may be due to the modification of transcription factor binding sites or the emergence of cis-regulatory elements leading to new TF expression profiles (Winkelmüller et al. 2021). Accordingly, shifts in gene regulation provide a more flexible means of adapting to changing environmental conditions and finetuning genomic predisposition (Koenig et al. 2013). The evolution of gene regulatory networks has been subject to intensive debates, with two main hypotheses proposed to explain regulatory diversity: hub conservation, where downstream differentiation occurs, versus hub divergence, where central regulatory genes shift in function. Some studies suggest that regulatory hub genes might remain conserved (Masalia et al. 2017). However, Wei et al. (2024) emphasised the importance of hub gene divergence and subsequent network topology shifts as the key driver of *S. chilense* adaptation to drought stress. Indeed, network re-wiring, potentially via modifying a single core gene, can significantly affect phenotypic plasticity (Koubkova-Yu et al. 2018). This may lead to highly specific responses, such as detoxification or enhanced protein localisation processes, as observed in this study. A recent study by Dong et al. (2024) found evidence for GRN-rewiring by reimplementing an ancient transcription factor, which altered the response to nutrient status in *Marchantia polymorpha.* Here, we present the first evidence of gene regulatory network rewiring leading to increased levels of QDR against a necrotrophic pathogen. Interestingly, both conserved and shared QDR-GRNs significantly enrich evolutionary younger genes dating back to Mesangiospermae-Solanales and discriminate older genes. Accordingly, we speculate that the adaptation to biotic stresses is modulated via shifts in the regulation and not adaptation via gene family founder events. Following this, core genes might be subject to negative selection, but regulating core TFs and the periphery might facilitate nuanced adaptation (Fagny and Austerlitz 2021).

### Co-optation of a TF might govern QDR in *S. pennellii* LA2963

Co-optation/exaptation or gene duplication followed by divergence are two potential mechanisms driving neofunctionalisation in plant-parasite interactions (Macquet et al. 2022; Plachetzki and Oakley 2007). Especially, the co-optation of transcription factors might display a major source of new regulatory patterns in GRNs (Artur and Kajala 2021). We identified a NAC29 transcription factor as a key modulator of QDR in *S. pennellii.* NAC transcription factors govern a wide array of developmental processes (e.g. flowering, cell cycle progression, and cell division) and may play a role in abiotic stress resistance (Zhang et al. 2018b; Nakashima et al. 2012; Kim et al. 2006; Ueda et al. 2020; Gonzalez-Bayon et al. 2019). Although some members of the NAP subgroup of NAC TFs might be involved in biotic stress response, only a few studies have linked them to resistance against necrotrophic pathogens (Lu et al. 2024; Tominello-Ramirez et al. 2024; Ma et al. 2021; Zhang et al. 2018a; Wang et al. 2020a; Masri and Kiss 2023). However, their role in promoting ROS tolerance could bolster disease resistance against *S. sclerotiorum* (Liu et al. 2014; Kabbage et al. 2015). The potential of NAC TFs in regulating diverse, species-specific processes highlights their evolutionary plasticity and adaptability. Interestingly, we observed differential regulation of NAC29 and changes in the downstream targets. However, the identification of co-optation events can be challenging.

By studying the phylotranscriptomics signature of resistance-associated GRNs, we can significantly enrich our understanding of QDR by integrating gene expression data with gene age (phylostratigraphy). By classifying genes into evolutionary “strata” and evaluating their expression profiles during pathogen challenge, we can assess whether older, highly conserved defence mechanisms or younger, lineage-specific expansions (perhaps newly co-opted for defence) drive quantitative resistance. We identified peaks in gene-family founder events that we can link to predicted genome expansions, such as a putative whole-genome triplication before solanales divergence. (Clark and Donoghue 2018; Huang et al. 2023; Maheepala et al. 2019). Subsequent expansions in angiosperms (Magnoliopsida) correlate with the reported emergence of flowering-specific genes, phytohormone signalling, pathogen defence, and secondary metabolism, underscoring genome duplication’s role in trait diversification (Huang et al. 2023; Maheepala et al. 2019; Clark and Donoghue 2018; Chanderbali et al. 2016; Bowles et al. 2020). Furthermore, the subsequent rise in taxonomically restricted genes within the Solanaceae may be attributed to the same genome triplication event, suggesting that such duplications drive species-specific adaptations and innovations (Clark and Donoghue 2018). Although we found evidence for species-specific gene-family expansion, we did not observe a link between taxonomically restricted genes on species level with QDR.

## Conclusion

Based on the phylotranscriptomic signature (i.e., shared copy numbers across *Solanum* species, broad basal expression patterns, and an ancient origin dating back to cellular organisms) and its unique functional role in *S. pennellii* LA2963, we propose that NAC29 has been co-opted for resistance-related functions against a necrotrophic pathogen. The parallel shifts in TF expression and downstream genes suggest that different mechanisms determine NAC29 co-optation. Possibly, altered TF regulation and differentiation of TF targets might underlie strong species-specific variability (e.g., cis-/trans-regulation and/or chromatin modifications, Jones and Vandepoele 2020). Possibly, the combination of trans- and cis-regulation might influence the equilibrium between conserved and specified QDR responses, as cis-regulation might underlie stronger differentiation (Wu et al. 2021). However, further research is needed to clarify the mechanisms underlying the co-optation of transcription factors driving resistance (Yuan et al. 2019). Additionally, it remains unclear how many ‘silent’ regulatory changes without a phenotypic effect (so-called developmental system drift) might influence the capacity to respond to environmental stress (Halfon 2017). Overall, NAC29 may serve as a prime example of how transcription factors can be repurposed for QDR, illustrating the broader role of TF co-optation in plant adaptation.

## Acknowledgements

We warmly thank Bettina Bastian for maintaining the plant material, Susanne Kleingarn for her wet lab support and Janina Fuß (CCGA) for her support during the sequencing. We also thank the TGRC at UC Davis (USA) for curating the germplasm used in this study. This work was partly funded by the DFG (STA1547/6) and the Agence Nationale pour la Recherche (ANR-21-813 CE20-30).

## Author Contributions

RS & SE designed the research, SE performed the experiments and conducted bioinformatic analysis, and SE & CTR & FD performed the data analysis. SE & RS wrote the manuscript. All authors read and approved the manuscript.

