## Supplemental Figures for "Co-optation of Transcription Factors Drives Evolution of Quantitative Disease Resistance Against a Necrotrophic Pathogen"

**Short title: Co-optation Drives QDR Against Necrotrophic Pathogens.^[1]^**

### Supplementary Figures

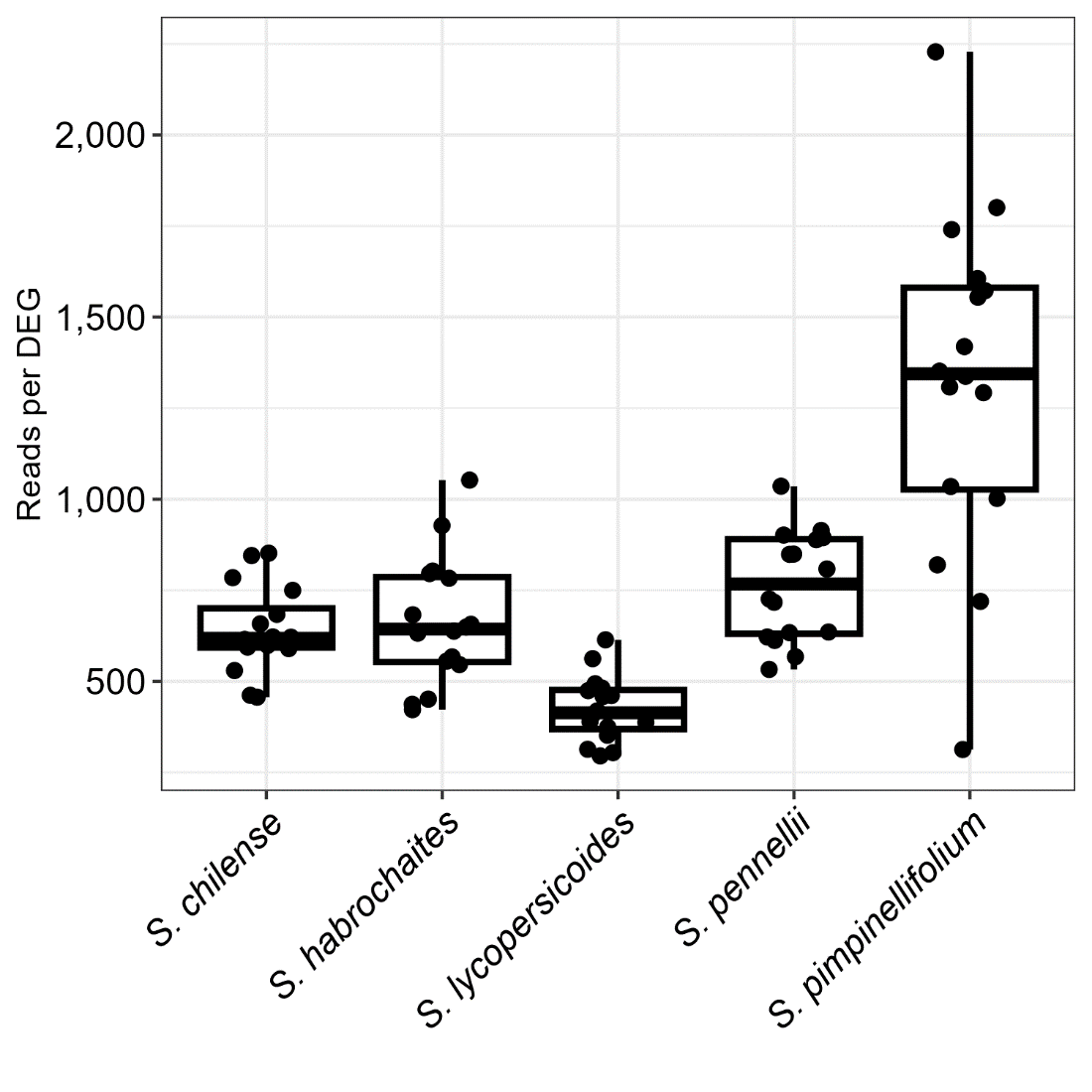

**Suppl. Figure 1: Relationship of read count and number of assigned DEGs.** We calculated the ratio based on the number of DEGs (contrast: infection vs. mock) per genotype and normalised it against the individual reads per sample (as represented by the dots).

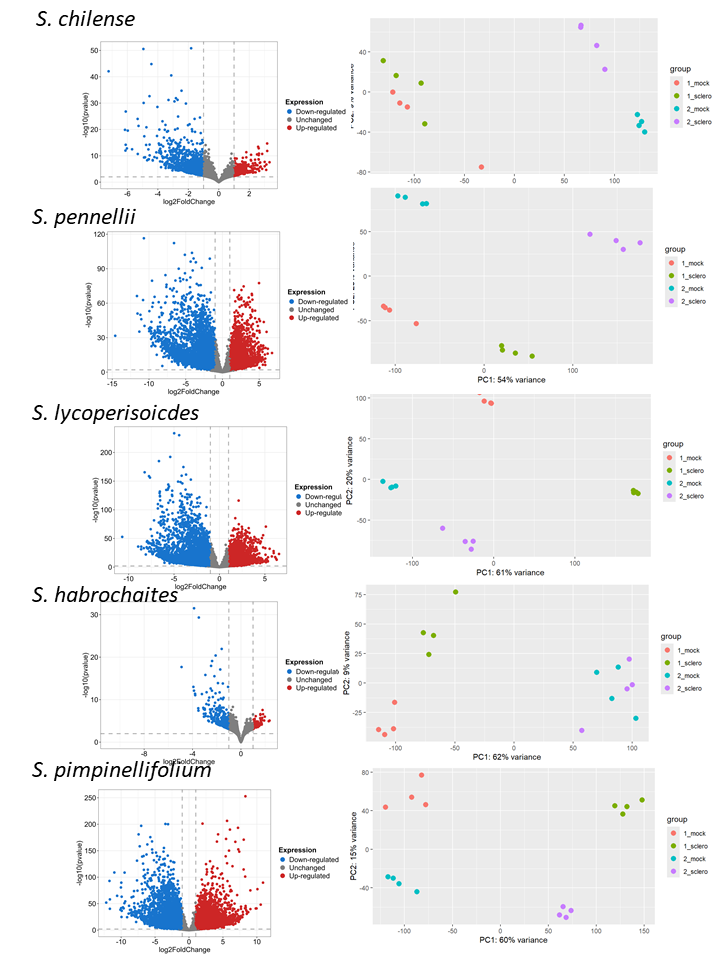

**Suppl. Figure 2:** **Basic Differential Gene Expression Analysis of all five tomato species.** The volcano plots show log2 foldchange against the pvalue of all five tomato species (contrast: Infected conditions res.-sus. genotype). Upregulated genes (red) are compared with downregulated genes (blue). The principal component analysis shows a clear separation of genotypes and treatments. Dots represent individual samples, with the colour defining the experimental treatment (1-susceptible, 2-resistant, mock and infected).

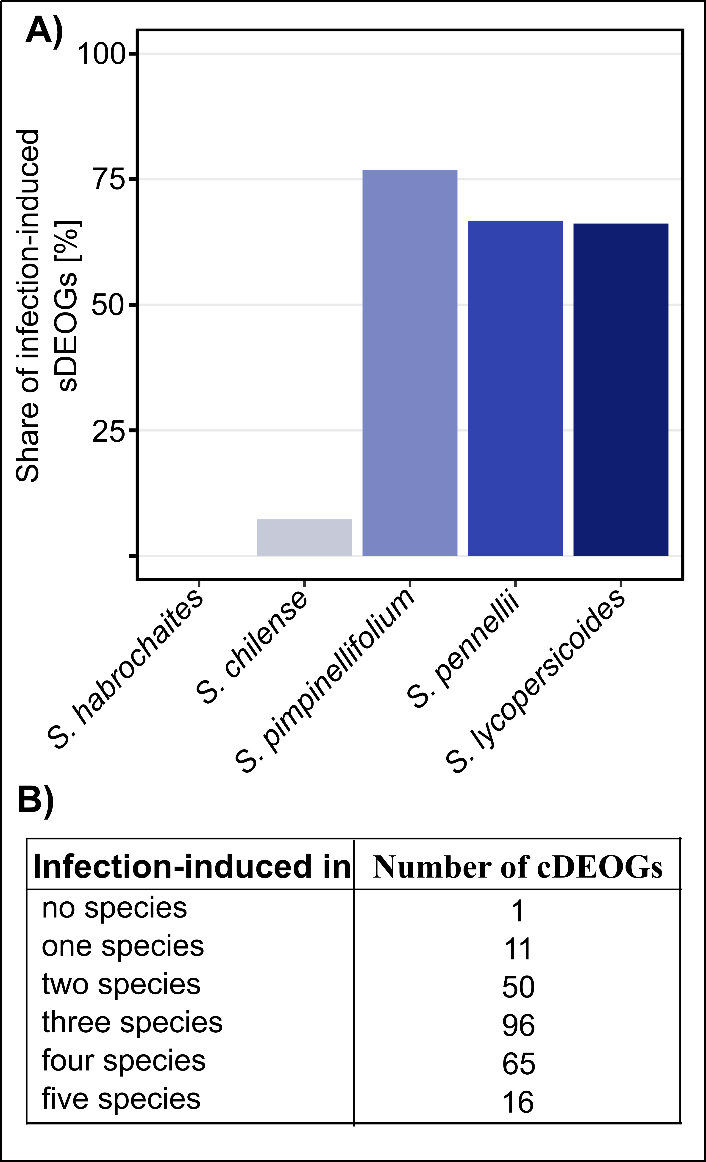

**Suppl. Figure 3: Amount of infection-induced differentially expressed orthogroups.
A)** species-specific sDEOGs and **B)** cDEOGs.

B)
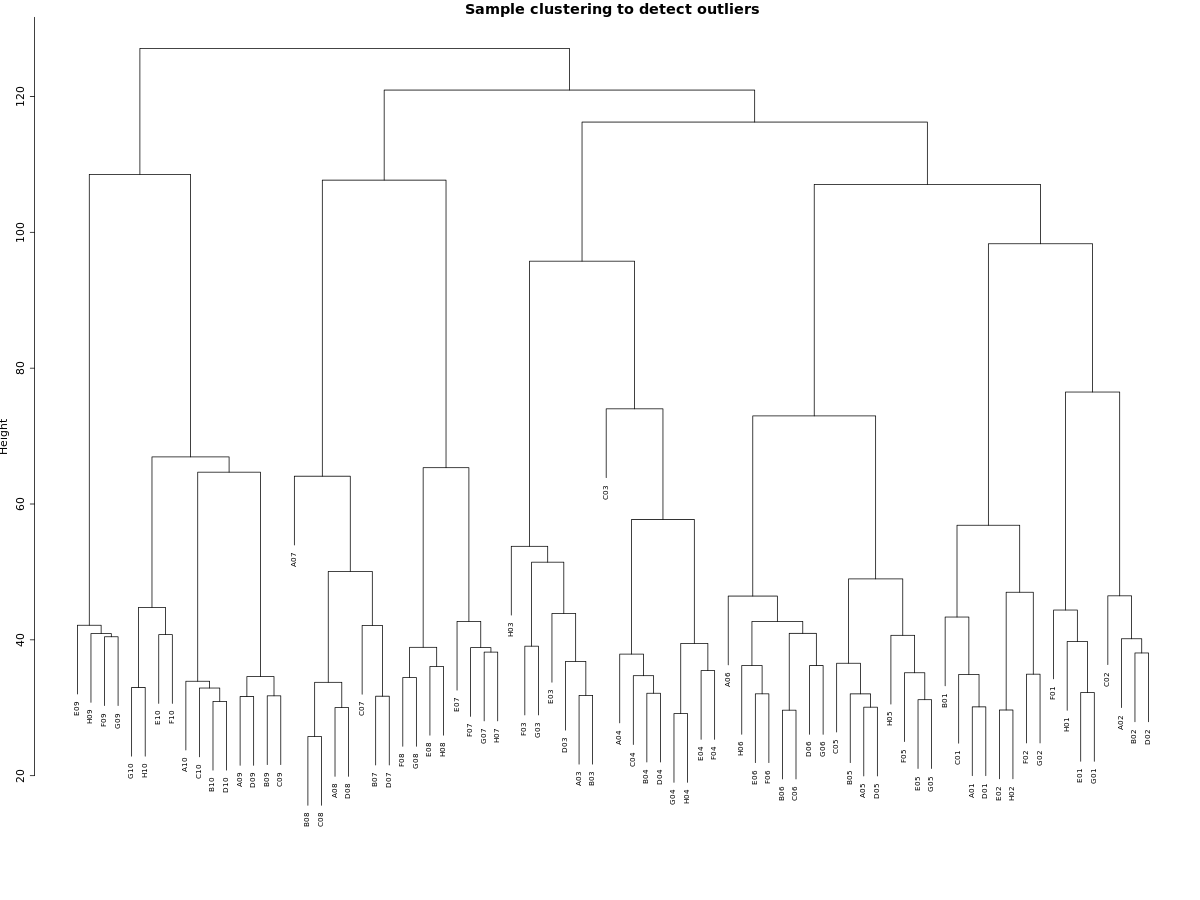

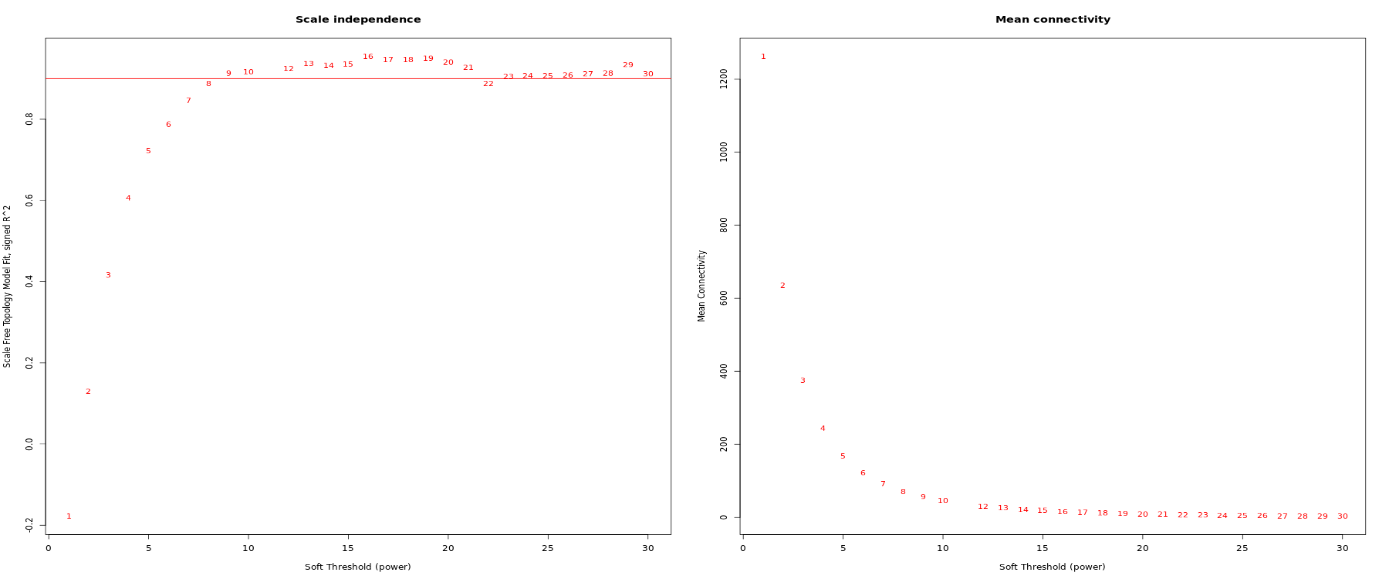

**Suppl. Figure 4: Sample evaluation for the OG-based WGCNA.
A)** Hierarchical clustering of all samples used in the study. **B)** Selection of the Soft power threshold for WGCNA network construction on the OG-data set. The red line indicates the signed R² threshold of 0.85. The SFT threshold was selected based on the scale-free topology model's plateau and mean connectivity.

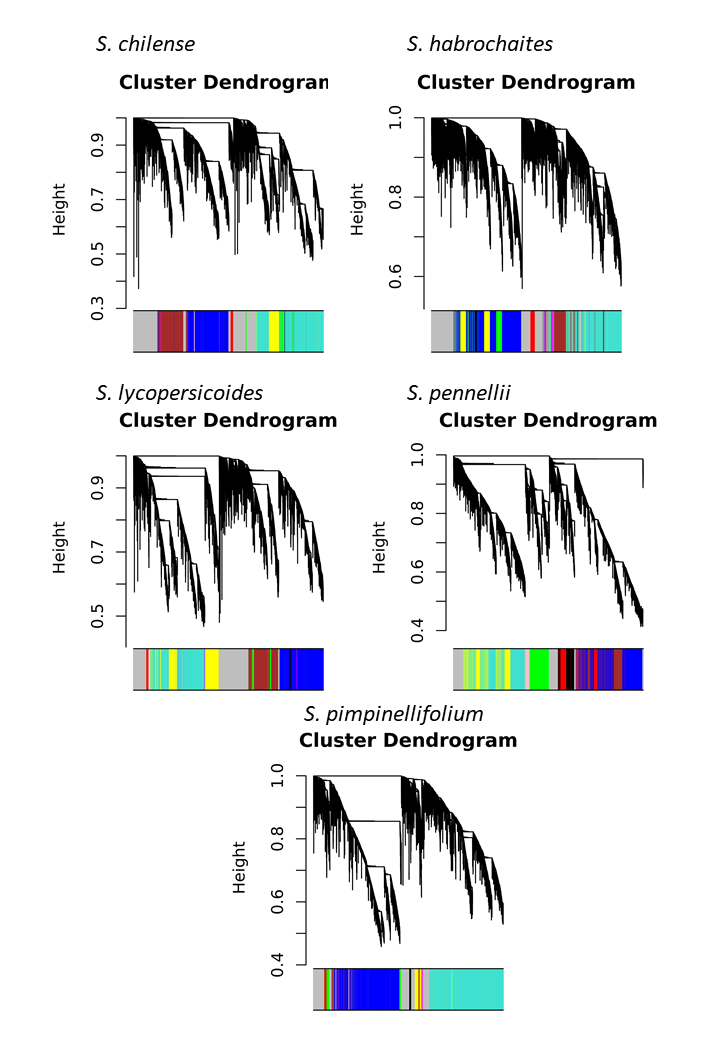

**Suppl. Figure 5: WGCNA dendrogram showing network topology and module assignment of each species.**

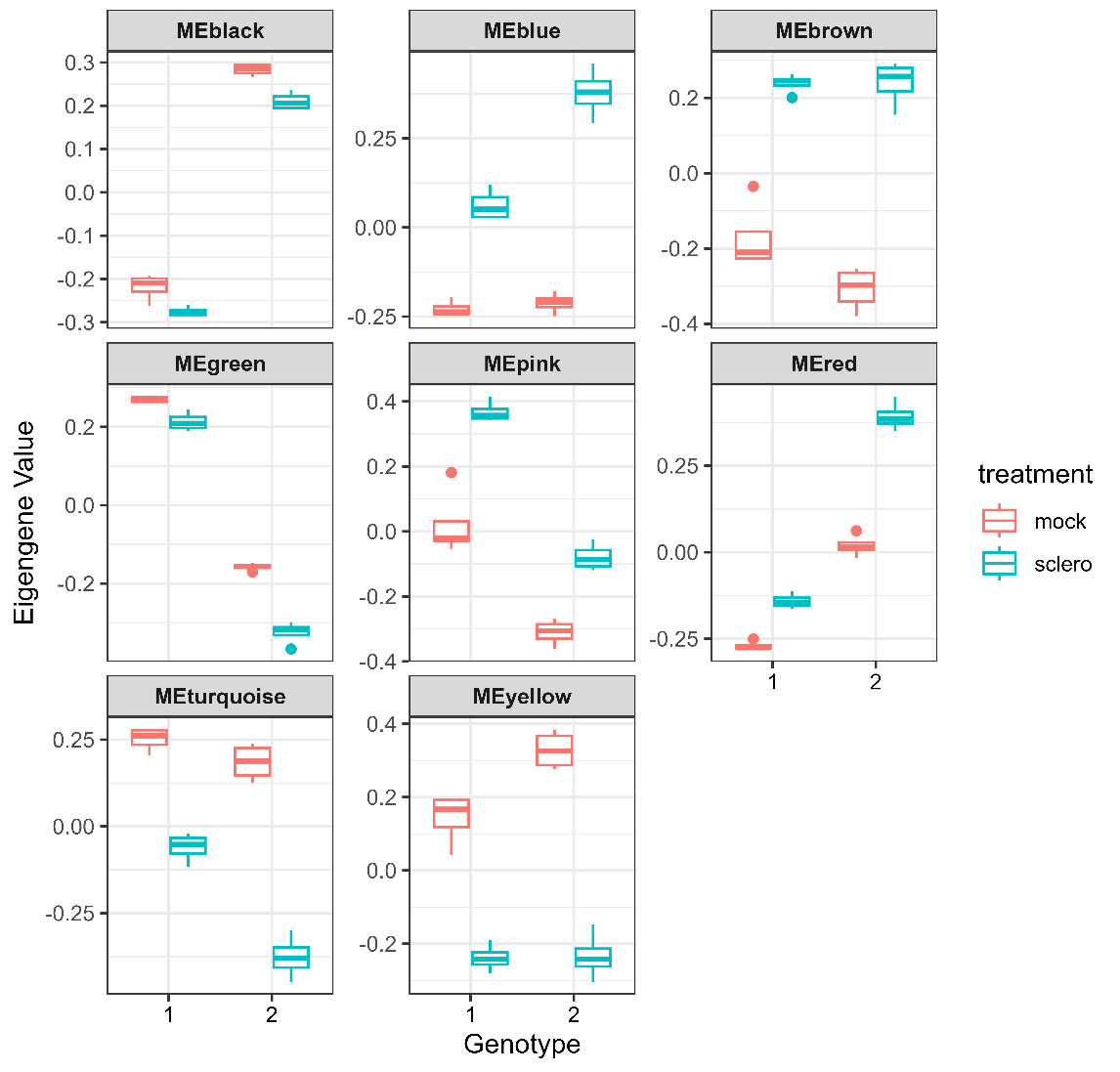

**Suppl. Figure 6: Module Eigengenes of the *S. pennellii* network.** The individual facets define coexpression modules assigned by the WGCNA. The Y-axis denominates the module eigengenes (MEs) of the two genotypes (1 susceptible, 2 resistant). The colour indicates experimental treatment. Boxplots summarise four independent samples.
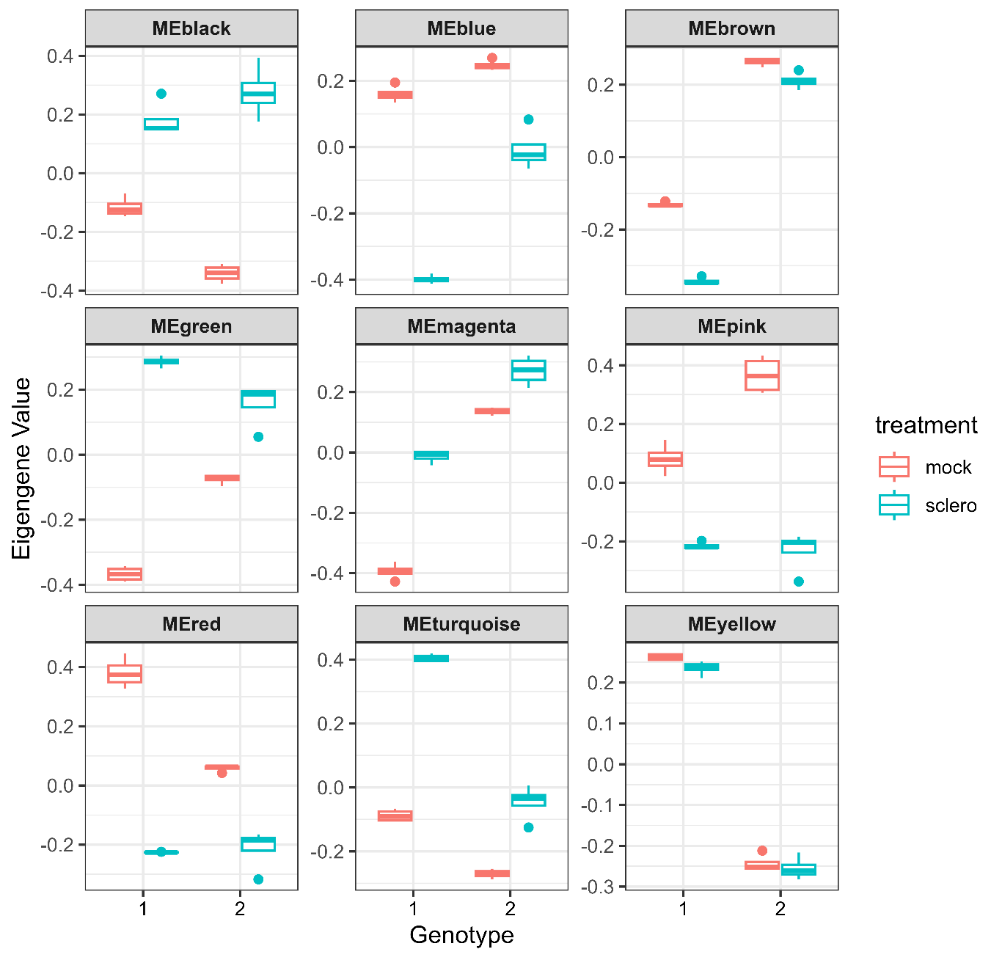

**Suppl. Figure 7: Module Eigengenes of the *S. lycopersicoides* Network.** The individual facets define coexpression modules assigned by the WGCNA. The Y-axis denominates the module eigengenes (MEs) of the two genotypes (1 susceptible, 2 resistant). The colour indicates experimental treatment. The boxplots summarise four independent samples.

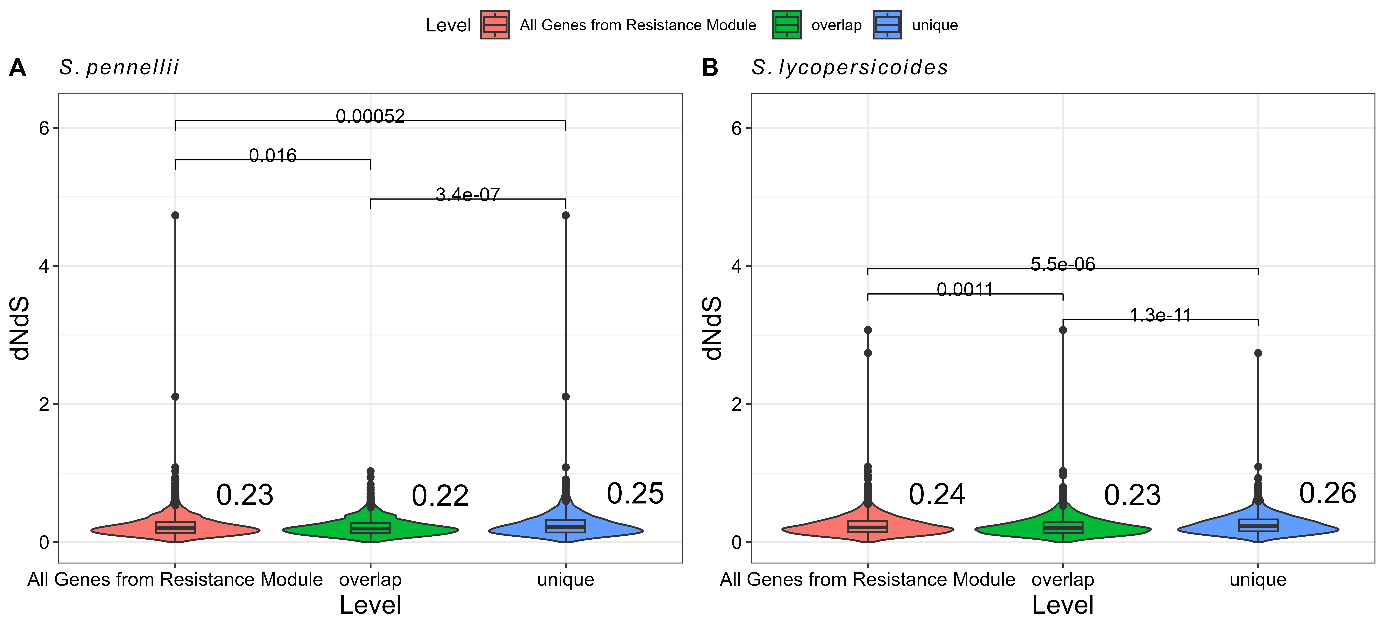

**Suppl. Figure 8: dNdS ratios contrasting overlapping vs. unique vs. all genes from the resistance modules on A) *S. pennellii* and B*) S. lycopersicoides*.** Statistical significance was determined using the Wilcoxon test. Numbers next to the violin charts represent the mean.

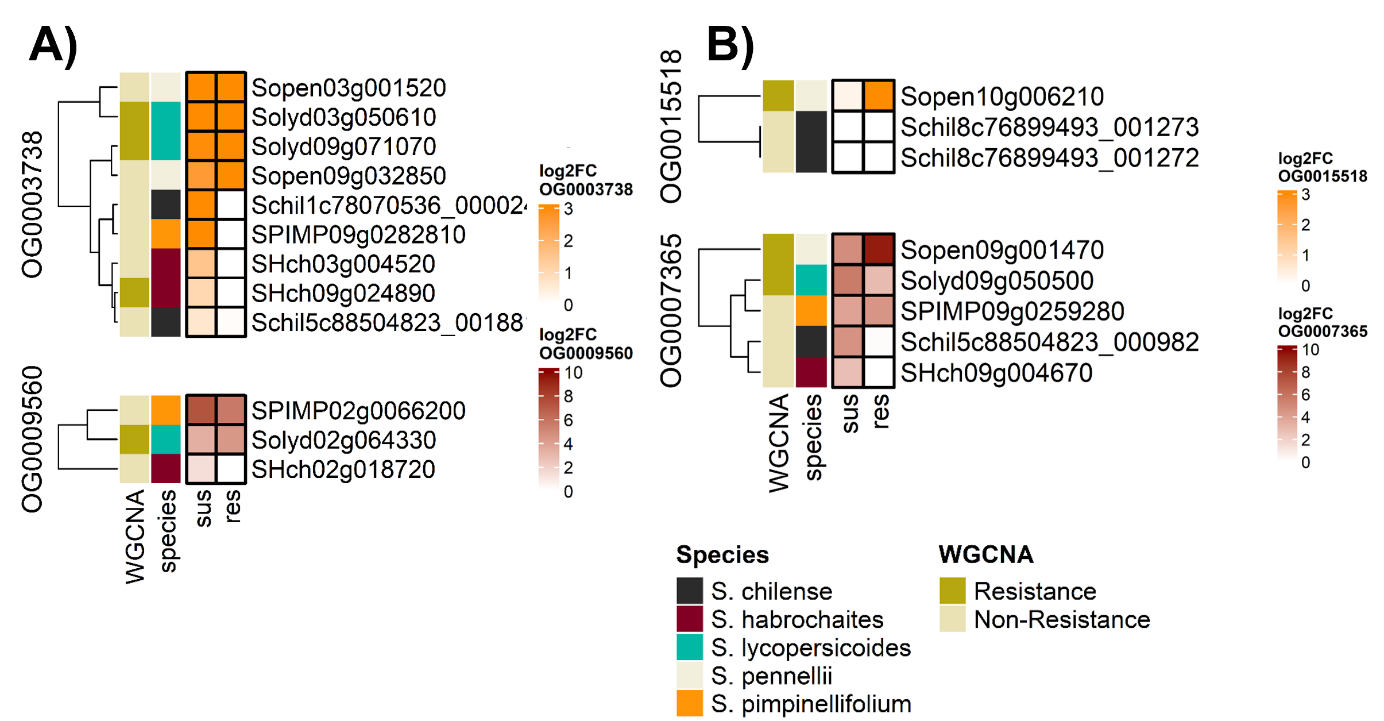

**Suppl. Figure 9: Regulatory pattern of resistance-associated TFs of A) *S. lycopersicoides* and B) *S. pennellii* compared across the other species using orthogroups.** The expression of each gene within the respective orthogroup was visualised across varying levels of QDR (susceptible | resistant), with colour coding indicating the respective species and the assignment of each TF to its species-specific resistance module

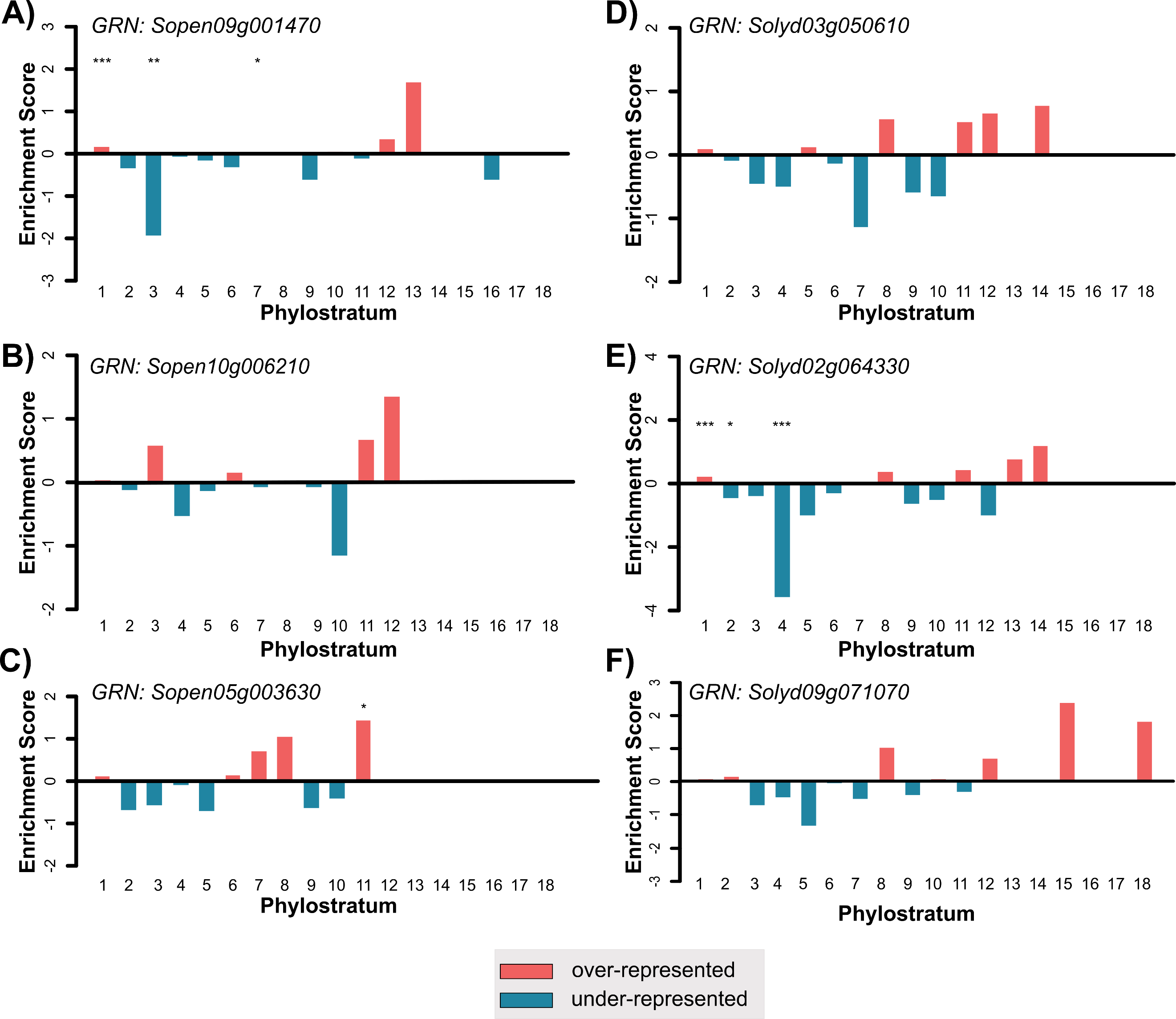

**Suppl. Figure 10: Phylostratum enrichment analysis per species of the three focal transcription factors.** We performed phylostratum enrichment analysis on the gene-regulatory networks downstream of three TFs on *S. pennellii* **(A-C)** and *S. lycopersicoides* **(C-E).** The enrichment score is indicated by the y-axis, and the respective phylostrata (cellular organisms till solanum) are located on the x-axis. Stars indicate the significance level of the respective enrichment after Fisher’s exact test and BH FDR correction.
